# Benchmarking Recent Computational Tools for DNA-binding Protein Identification

**DOI:** 10.1101/2024.09.01.610735

**Authors:** Xizi Luo, Amadeus Song Yi Chi, Andre Huikai Lin, Tze Jet Ong, Limsoon Wong, Chowdhury Rafeed Rahman

## Abstract

Identification of DNA-binding proteins (DBPs) is a crucial task in genome annotation, as it aids in understanding gene regulation, DNA replication, transcriptional control and various cellular processes. In this paper, we conduct an unbiased benchmarking of eleven state-of-the-art computational tools as well as traditional tools such as ScanProsite, BLAST, and HMMER for identifying DBPs. We highlight the data leakage issue in conventional datasets leading to inflated performance. We introduce new evaluation datasets to support further development. Through a comprehensive evaluation pipeline, we identify potential limitations in models, feature extraction techniques and training methods; and recommend solutions regarding these issues. We show that combining the predictions of the two best computational tools with BLAST based prediction significantly enhances DBP identification capability. We provide this consensus method as user-friendly software. The datasets and software are available at: https://github.com/Rafeed-bot/DNA_BP_Benchmarking.

**Key Points:**

- We designed a comprehensive evaluation pipeline which systematically evaluates eleven recent machine learning (ML) based DBP identification tools.
- We analyzed the test prediction mistakes made by top-performing tools identifying their potential limitations in terms of model architecture, feature extraction and class balancing.
- We showed that although the best of these tools do not convincingly outperform BLAST, they still provide substantial value when integrated together with BLAST into a simple majority-voting ensemble.
- We provide recommendations on more robust development & evaluation and better usability of future tools.
- We provide the two best-performing ML-based tools, BLAST and the ensemble method as user-friendly software, as well as our proposed datasets, publicly available via GitHub.

## 2. Introduction

Identification of DNA-binding proteins (DBP) is a fundamental task in molecular biology. DBPs have a wide range of significant applications including the development of drugs, antibiotics and steroids for various biological effects. They are also extensively used in biophysical, biochemical and biological studies of DNA [1]. DBPs bind to DNA through a variety of mechanisms, often mediated by specific structural motifs that recognize and interact with particular DNA sequences or structures. These interactions are typically governed by a combination of hydrogen bonds, van der Waals forces and ionic interactions between the protein’s amino acid residues and the DNA’s nucleotide bases or phosphate backbone. One of the most well-known motifs is the helix-turn-helix (HTH) motif commonly found in transcription factors. The HTH motif consists of two α-helices separated by a short turn. One helix fits into the major groove of the DNA allowing specific base pair recognition while the other stabilizes the interaction [2, 3]. Another common motif is the zinc finger where a zinc ion stabilizes a loop of amino acids that interacts with the DNA [4]. Additionally, proteins can bind to DNA in a sequence-specific manner or in a sequence-independent manner. Sequence-specific interactions involve proteins recognizing particular nucleotide sequences such as the binding of transcription factors to promoter regions to regulate gene expression [5]. Conversely, sequence-independent interactions involve proteins binding to DNA based on its structure or overall shape such as the binding of histones to form nucleosomes [6].

Given the importance of DBPs, various experimental methods have been employed to identify them such as filter binding assays [7], genetic analysis [8] and chromatin immunoprecipitation on microarrays [9]. However, these experimental methods are time-consuming and expensive [10]. Computational tools for DBP identification are essential due to the vast amount of genomic data generated by high-throughput sequencing technologies [11]. Our motivation for this benchmarking study stems from the need to systematically evaluate and compare the performance of existing computational tools for DBP identification. With many tools available, each employing different models, datasets, and features, there is a lack of systematic study regarding their effectiveness. Another goal of this study is to evaluate the performance gain of these tools over simple BLAST search to understand the necessity of these tools for DBP identification.

Machine learning-based computational methods have gained prominence in recent years for DBP identification leveraging advances in statistical techniques and feature engineering. These methods can be broadly categorized based on the models they employ and the features they utilize for prediction. Traditional machine learning (ML) models such as Support Vector Machines (SVM) [12, 13, 14, 15, 16], Random Forests (RF) [17, 18, 19], and nearest neighbors algorithm [20] have been used for DBP prediction. These models typically rely on manually curated features extracted from protein sequences and structures. More recent approaches employ deep learning architectures such as Convolutional Neural Networks (CNN) [21, 22], Recurrent Neural Networks (RNN) [22, 23, 24] and pretrained protein large language models (as feature extractors) [25, 26]. CNNs are adept at capturing spatial hierarchies in protein sequences, while RNNs effectively model sequential dependencies. PLMs on the other hand relies on self attention mechanism [27] (similar to Transformer architecture) in order to have a deeper understanding on relative importance among amino acids of a protein sequence. Although these models often require large datasets for training and fine-tuning, they have reported superior performance in capturing the intricate characteristics of DNA-binding sites.

Features used for prediction can be derived from various sources. Sequence-based features are obtained directly from the amino acid sequences of proteins. Common sequence-based features include amino acid composition [28, 29], dipeptide composition [30, 31], physicochemical properties such as hydrophobicity and charge [12, 32, 33], and PLM embedding [34, 35]. Given the importance of the three-dimensional conformation of proteins in DNA binding, structural features provide valuable insights. These include secondary structure elements (e.g., alpha helices, beta sheets), solvent accessibility [36], and binding site geometries. Position-specific scoring matrices (PSSMs) and profiles derived from multiple sequence alignments capture evolutionary conservation signifying functional importance which can aid in identifying potential DNA-binding regions [14, 37, 38, 34].

One major issue with these methods is the quality and representativeness of the datasets used for training and evaluation. Existing datasets suffer from bias and data leakage leading to inflated performance. Another issue is that some tools primarily focus on various feature extraction techniques from the protein sequences which may include irrelevant or inappropriate feature sets not targeted towards DBP prediction resulting in overfitting. Additionally, the models employed have bias towards specific characteristics of the protein sequences which causes mispredictions.

In this paper, we introduce novel benchmarking datasets which are designed to provide a more realistic and unbiased evaluation of DBP prediction tools. We benchmark eleven state-of-the-art tools analyzing their models, datasets, and feature types and identify the top-performing tools through rigorous evaluation. Furthermore, we explore the potential of traditional tools like BLAST [39], ScanProsite [40], and HMMER [41] for DBP prediction providing insights into their capabilities and limitations. We show that the two best computational tools (according to our evaluation) ensembled with BLAST significantly improves performance. Finally, we analyze the common mistakes made by the top-performing tools to summarize potential causes and possible solutions. We provide our curated train-test split and the ensemble method as user-friendly software via publicly accessible GitHub repository.

## 3. Benchmarked Tools

The benchmarked tools can be categorized based on the models they proposed, the feature representation methods they employed and the datasets they utilized (Table 1 and Supplementary Data 8). We discuss each categorization in this section. We conclude this section by discussing the issues we faced in running these eleven recent computational tools.

**Table 1.** Benchmarked tool summary.

| Tool (Publication Date) | Feature Type | Computation Model | Dataset (Train + Test) | Web Server (YES / NO) | Code Availability |
| --- | --- | --- | --- | --- | --- |
| Local-DPP [42]<br>(23/06/2016) | Evolutionary | Classic | PDB1075 + PDB186 | NO | Not available |
| DNABP [43]<br>(01/12/2016) | Sequence + Evolutionary | Classic | PDB14K (97%) + PDB14K (3%) | NO | Not available |
| iDNAProt-ES [37]<br>(02/11/2017) | Sequence + Evolutionary + Structure | Classic | PDB1075 + PDB186 | YES | Not available |
| StackDPPred [44]<br>(19/07/2018) | Evolutionary + Structure | Classic (stage) | PDB1075 + PDB186 | YES | Available but not runnable |
| PseAAC [45]<br>(11/10/2018) | Sequence | Classic | PDB1075 + PDB186 | NO | Available |
| DeepDBP [46]<br>(19/03/2020) | Sequence | Deep Learning | PDB1075 + PDB186 | NO | Partially available (no feature extraction) |
| PDBP-Fusion [24]<br>(03/05/2021) | Sequence | Deep Learning | PDB14K + PDB2272 | NO | Not available |
| KK-DBP [47]<br>(29/11/2021) | Evolutionary | Classic | PDB1075 + PDB186 | NO | Not available |
| LSTM-CNN_Fusion [23]<br>(02/06/2022) | Sequence + Evolutionary | Deep Learning | PDB14K + PDB2272 | NO | Available |
| PB_DBP [35]<br>(04/12/2023) | Sequence | Deep Learning | A custom dataset (80% + 20%) | NO | Not available |
| PreDBP-PLMs [34]<br>(08/07/2024) | Sequence + Evolutionary | Deep Learning | PDB1075 + PDB186<br>PDB14K + PDB2272 | NO | Available |

### 3.1 Feature Representation Based Categorization

The feature representation methods can be categorized into three types: sequence, evolutionary and structure-based feature representation (summarized in Table 1). Sequence-based features are derived from the original amino acid sequences. Common techniques include one-hot encoding, physicochemical properties, amino acid composition and others. Starting with the most basic approach, PDBP-Fusion and LSTM-CNN_Fusion used one-hot encoding on the sequence residues, allowing the classifiers to learn features automatically. More Recent tools such as PB_DBP and PreDBP-PLMs encoded DNA sequences through pretrained protein language models (PLM). In contrast, PseAAC and DeepDBP examined the frequency of single amino acids, dipeptides, and tripeptides, as well as the distribution of single amino acids. They also focused on the frequency of non-consecutive amino acids and their distributions. DNABP and iDNAProt-ES both utilized physicochemical property features as part of their final feature vectors. These features categorize each amino acid residue based on properties such as hydrophobicity, polarity, and polarizability; and then calculate the composition, transition, and distribution of these groupings [43, 37]. Additionally, DNABP included a sequence-based feature generated by DNABR [48], a tool that predicts DNA-binding residues and their corresponding reliability indices given a sequence. Sequence-derived features have the advantage of being straightforward to compute and do not require evolutionary information, making them faster and more suitable for large-scale analyses. However, since protein sequence is often not a good representative of its 3D secondary structure [49, 50], relying solely on sequence-based features may not capture the complex patterns associated with DBPs.

Evolutionary features are derived from position-specific scoring matrices (PSSMs) generated by PSI-BLAST [51]. For instance, Local-DPP and iDNAProt-ES utilized the average probability of each residue position mutating to one of the 20 residue types during the evolutionary process. Unique to Local-DPP is its consideration of local features by segmenting the PSSM into several sub-PSSMs of the same size before applying any feature extraction technique. Another commonly used feature set involved selecting rows in the PSSM corresponding to the same amino acid type and summing the values in each column (used by DNABP, StackDPPred, KK-DBP, and PreDBP-PLMs). In DNABP, these 20 values were combined with physicochemical property values through some arithmetic operations. Additionally, one feature set involved the occurrence probabilities for pairs of the same amino acids separated by a certain distance along the sequence, while another feature set involved pairs of different amino acids; both calculated from PSSM matrix [37, 44, 47, 34]. It is noteworthy that KK-DBP not only utilized conventional PSSMs with 20 columns but also included features generated by reduced PSSMs (RPSSMs) calculated as the average square of the sum of PSSM values from different columns grouped at specific distances. KK-DBP and PreDBP-PLMs also proposed column-wise variances derived from the reduced PSSM as one of its final feature sets. Unique to iDNAProt-ES, this tool proposed a feature set that considered the column-wise distribution of values in the PSSM retrieved by calculating partial sums column-wise. Finally, as a deep learning-based tool, LSTM-CNN_Fusion used PSSMs as direct input to the CNN model for automatic feature extraction without employing any PSSM transformation. Evolutionary features derived from PSSMs have proven to be highly informative for DBP prediction, as demonstrated by the superior performance of tools like Local-DPP and LSTM-CNN_Fusion (discussed later). These features capture evolutionary conservation patterns that are often associated with functional regions in proteins. However, processing PSSMs can be challenging due to their variable length (corresponding to protein sequence length). Methods like truncating the PSSM matrix to a fixed size or deriving fixed size summary statistics feature vector may lead to significant loss of information, especially for longer sequences. Additionally, the reliance on PSI-BLAST for PSSM generation can be computationally intensive.

For structure-based feature representation, the three-dimensional geometry of proteins is leveraged to identify potential DBPs. iDNAProt-ES proposed a set of structural features extracted from information provided by SPIDER2 [52] as an SPD file. These features included the occurrence and composition of secondary structures, accessible surface area (ASA), torsional angles and structural probabilities. Secondary structure occurrence calculated the frequency of three types of structural motifs in proteins: α-helix (H), β-sheet (E), and random coil (C). Structural probabilities represented the likelihood of each amino acid being in H, E, or C conformation. Beyond these basic features, the matrix in the SPD file was processed similarly to PSSM-related features. For instance, the average products of values from both the same and different columns grouped at specific distances were calculated for torsional angles and structural probabilities. StackDPPred proposed integrating the Residue-wise Contact Energy Matrix (RCEM) [53] into its feature set. The RCEM feature (20 *×* 20 matrix) approximates the structural stability of proteins by using predicted residue contact energies derived from known 3D structures in order to account for amino acid interactions and intrinsically disordered regions (IDRs) in proteins. Structure-based features can provide insights into the spatial arrangement and folding patterns of proteins, which are crucial for DNA-binding activity. However, obtaining accurate structural information can be resource-intensive and time-consuming. Additionally, the inclusion of numerous structural features derived from predicted models may introduce errors and increase the feature dimensionality significantly, leading to overfitting [54]. Incorporating structural features requires careful consideration to balance the benefit of additional information against the risk of model complexity and data quality.

### 3.2 Computational Model Based Categorization

We discuss two distinct groups of modeling -one group utilized classic ML models that relied on complex feature extraction techniques, while the other group employed deep learning models capable of automatic feature extraction and end-to-end training. The most commonly used classic ML model was the RF Classifier [55] used by Local-DPP, DNABP, StackDPPred, PseAAC and KK-DBP. iDNAProt-ES exclusively used Support Vector Machine (SVM) [56], while PseAAC used Extra Tree Classifier. StackDPPred proposed a stacking framework which involved two stages of learning. The base classifiers included SVM, Logistic Regression [57], KNN [58] and RF, while the meta-classifier was another SVM with a radial basis function (RBF) kernel [59]. Classic ML models like Random Forest and SVM offer advantages such as interpretability, faster training, and robustness with small datasets. However, they require fixed-size feature vectors, which can lead to information loss for variable-length sequences. Handling heterogeneous features can also introduce noise if not managed carefully. While simpler than deep learning models, they may struggle with complex sequence patterns and dependencies.

In the deep learning group, a combination of CNN [60] and Long Short-Term Memory networks (LSTM) [61] was utilized by both PDBP-Fusion and LSTM-CNN_Fusion. Unlike PDBP-Fusion, which stacked the LSTM layer on top of the CNN layers, LSTM-CNN_Fusion used these two models in parallel, with CNN extracting features from the PSSMs and LSTM learning information from the original protein sequences. DeepDBP experimented with both CNN and Artificial Neural Network (ANN) model. DeepDBP-ANN utilized sequence composition based summary statistics while DeepDBP-CNN automatically learned features from the raw sequence with the assistance of a trainable embedding layer and a convolution layer. Regarding the two PLM-related tools, PB_DBP employed a BiLSTM to process the output of a pretrained PLM model, while PreDBP-PLMs utilized a CNN for this purpose. Deep learning models offer several advantages, such as the ability to automatically learn complex patterns from raw data, making them well-suited for tasks involving sequence dependencies and large datasets. Architectures like CNNs excel at capturing local patterns, while LSTMs are effective at modeling sequential dependencies. However, these models come with drawbacks including high computational costs and the need for large datasets to prevent overfitting. Additionally, deep learning models are often less interpretable than traditional ML models, making it challenging to understand the underlying decision-making process. Finally, pretrained PLM embeddings can be biased based on the category of protein sequences they were mostly pretrained on.

### 3.3. Dataset Based Categorization

The tools can be divided into two groups based on the training dataset they used (column 4 of Table 1). DNABP, PDBP-Fusion, LSTM-CNN_Fusion, and PreDBP-PLMs all used PDB14K [43] as their training dataset. Among these tools, PDBP-Fusion, LSTM-CNN_Fusion and PreDBP-PLMs utilized PDB2272 [24] as their test dataset. DNABP took out 203 positive and 203 negative sequences from PDB14K and used them for independent testing. The remaining seven tools except PB_DBP used PDB1075 as the training dataset and PDB186 [36] as the test dataset. PB_DBP created a large dataset consisting of around 42K DBPs and non-DBPs, and split the dataset into 80% train and 20% test set. But this dataset has not been made publicly available. Details of the datasets used will be discussed in Section 4.

### 3.4. Tool Benchmarking Issues

None of the computational tools that we benchmarked had any usable software or web server available except for iDNAProt-ES and StackDPPred. Although iDNAProt-ES has its own web server, it is not effective as it requires PSSM evolutionary feature and SPD structural feature files and cannot work with raw sequences as input. The web server provided by StackDPPred is not accessible from Singapore. On top of that, none of the tools provided the trained models that could directly be used for prediction. As a result, we looked into their provided codebase to train and test these tools ourselves. Six out of the eleven tools did not have any code available and we could not retrieve the source codes even after corresponding with their authors (see the last column of Table 1). StackDPPred and DeepDBP had end-to-end running issues which required our attention. The codes provided by PseAAC, LSTM-CNN_Fusion, and PreDBP-PLMs were runnable. As mentioned in Supplementary Data 1, some of the tools had some inconsistencies between the provided code and the paper. In such cases, we simply followed the code. In some other cases, the selected features were not clearly indicated and as a result, we had to rerun the feature selection process. Furthermore, two feature values of DNABP required running of the DNABR classifier (predicts DNA-binding residue confidence scores) on the protein sequence of interest. Since the server of this classifier is currently not available, we had to run DNABP without using these two feature values. Finally, we attempted to achieve similar results as reported in the corresponding tool papers after training on the mentioned training set and testing on the mentioned test set, which would ensure correct replicability of the tools. Unfortunately, our achieved performance was much worse than the reported performance for two out of the eleven tools - iDNAProt-ES (around 40% drop in MCC score) and DeepDBP (over 80% drop in MCC score) marked in yellow in Supplementary Data 4. Furthermore, since the training and test datasets of PB_DBP were not available, we could not make sure that it’s reported performance was reproducible.

## 4. Current Dataset Details and Limitations

Among the eleven tools we benchmarked, two combinations of training and test datasets were commonly found - (1) PDB1075 as training & PDB186 as test, (2) PDB14K as training & PDB2272 as test. Table 2 contains the basic information of the four datasets. PDB1075 and PDB186 were originally curated by Liu et al. [62] and Lou et al. [36]. DBPs were initially acquired from the Protein Data Bank (PDB) [63] by mmCIF keyword of “DNA binding protein” through the advanced search interface. To mitigate fragmentary sequences, proteins with fewer than 50 and 60 amino acids were excluded from PDB1075 and PDB186, respectively. Additionally, proteins containing the residue ‘X’ were omitted to avoid unknown residues. Subsequently, PISCES [64] and NCBI’s BLASTCLUST [51] were employed to eliminate proteins exhibiting more than 25% identity with any protein within PDB1075 and PDB186, respectively. Similarly, the non-DBPs were randomly selected from other proteins in PDB and filtered using the same criteria.

**Table 2.** Commonly used dataset summary.

| Dataset | Source | +ve & -ve Sequence Number | Max, Min, Median Sequence Length | Data Curation Process |
| --- | --- | --- | --- | --- |
| PDB1075 | Liu et al. [62] | 525 & 550 | 1323, 50, 189 | Remove seqs $\geq 25\%$ similarity and irregular chars |
| PDB186 | Lou et al. [36] | 93 & 93 | 1323, 64, 208 | Remove seqs $\geq 25\%$ similarity and irregular chars |
| PDB14K | DNABP [43] | 7131 & 7131 | 4911, 47, 327 | Remove seqs $\geq 40\%$ similarity and irregular chars |
| PDB2272 | PDBP-Fusion [24] | 1153 & 1119 | 5184, 51, 325 | Remove seqs $\geq 25\%$ similarity and irregular chars |

In case of PDB14K, “DNA binding” was used as a keyword to search the UniProt database to obtain DBPs. Only sequences with lengths between 50 and 6000 amino acids were retained, and redundant data were removed using a 40% similarity threshold. To obtain the non-DBPs, all proteins from the UniProt database that lacked any implied RNA/DNA-binding functionality were obtained following a procedure proposed by Cai and Lin [12]. PDB2272 was curated in a similar fashion. Random selection was performed on the non-DBPs in order to make all these four datasets have almost equal number of DBPs and non-DBPs.

The primary limitations of these datasets can be categorized into three main points. Firstly, non-DBPs (negative) vastly outnumber DBPs (positive) in the real world. Consequently, datasets with approximately equal number of positive and negative samples do not accurately represent real-life scenario. The random undersampling of non-DBPs during training for class balancing eliminates important data points (these data maybe important for a comprehensive understanding of the full feature space of negative samples) potentially leading to a less robust model. The most alarming aspect is that this random undersampling was also performed in the test sets of these earlier works for class balancing; this would confound the extrapolation to real-life performance [65, 66].

Secondly, data leakage is prevalent between the training and test datasets described above. When CD-HIT [67] is applied at 40% similarity threshold, 85 out of the 93 DBPs in PDB186 test set exhibit >40% similarity to one or more DBPs in the corresponding training dataset PDB1075. Similarly, 1007 out of 1153 DBPs in PDB2272 test set exhibit >40% similarity to one or more DBPs in the corresponding training dataset PDB14K. Wang et al. [68] described the occurrence of highly similar data across training and test datasets as “data doppelgangers”. These high numbers of data doppelgangers result in models being repeatedly tested on data that are highly similar to what they encountered during training, leading to inflated performance. The fact that ML models have a tendency of overfitting to sequence type training data makes matters even worse [69]. The primary and secondary structure of proteins belonging to the same species can be quite different depending on their biological function [70] further emphasizing the need for non-redundant test data with respect to the training set.

Finally, sequences appearing in both the positive and negative sets cause ambiguity during training. PDB14K contains 74 sequences labeled as both positive and negative, while one sequence labeled as positive in PDB1075 appears as negative in PDB14K.

These limitations emphasizes the necessity for a new benchmarking dataset - one with minimized similarity between training and test datasets to more accurately evaluate the performance of various models under conditions where negative samples outnumber positive ones with no ambiguity.

## 5. Benchmarking Dataset Development

To address the shortcomings of current datasets, we constructed a new dataset named BTD. To construct the positive set, we first extracted DBPs from UniProtKB [71] selecting only those manually annotated with a review score of 5 (top subfigure of Figure 1). Sequences containing non-amino acid characters including ambiguous ones and those with lengths outside the range of 50 to 3000 amino acids were excluded. CD-HIT was applied with a similarity threshold of 40% to reduce redundancy within the extracted DBPs. To further prevent redundancy between training and test DBPs, we filtered out sequences similar to those found in commonly used training dataset (PDB14K and PDB1075) DBPs at 40% similarity threshold. This filtering step ensures the elimination of potential data leakage between the training datasets and BTD. Without this precaution, the test data in BTD would maintain internal non-redundancy but might still overlap with the training datasets, compromising the integrity of the evaluation process. The output refined dataset constitutes the positive component of BTD containing 16,384 sequences.

**Fig. 1.**
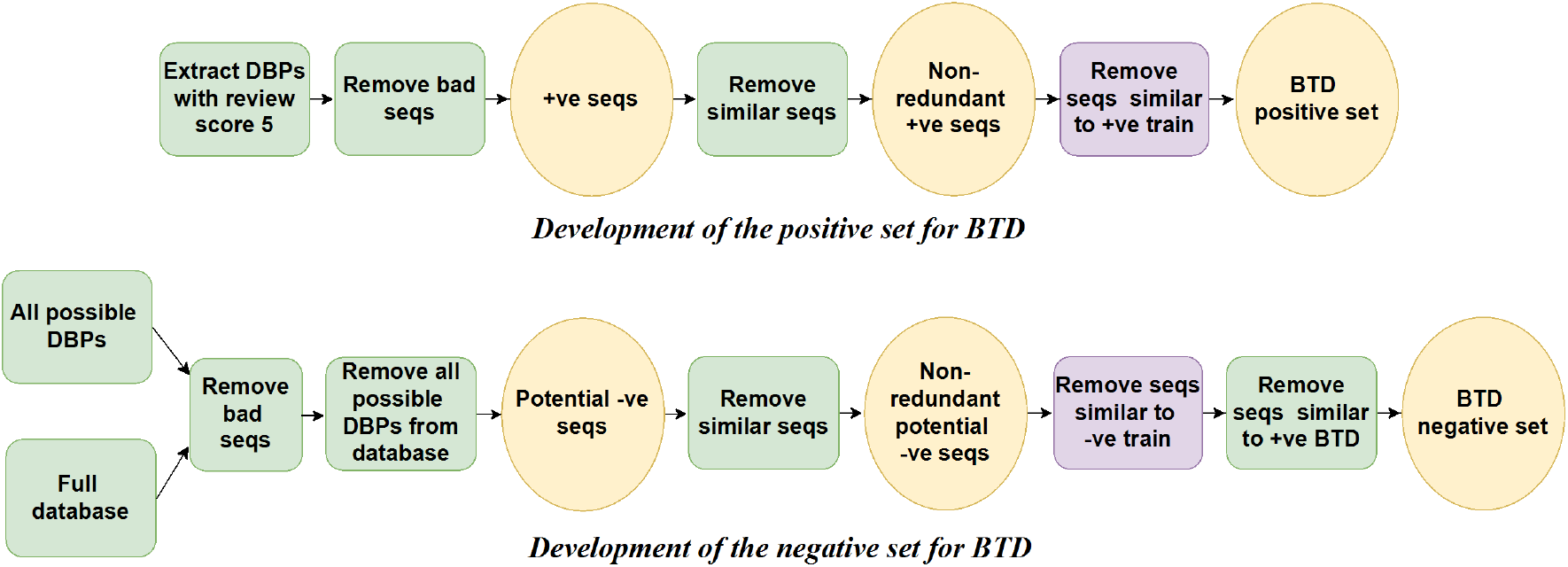
Benchmarking dataset BTD development pipeline.

To compile a negative set (bottom subfigure of Figure 1), we first identified all possible DBPs from UniProtKB by filtering with keywords indicative of RNA/DNA-binding functionality as described in [12]. Concurrently, the full Swiss-Prot database was also downloaded. The set of non-DBPs was obtained by excluding the possible DBPs from the Swiss-Prot database. Note that sequences containing non-amino acid characters and outside the length range of 50 to 3000 amino acids were not included. CD-HIT was then applied to the potential non-DBPs to remove redundancy with 40% similarity threshold. To avoid data doppelgangers between training and test sets, we further filtered out sequences similar to those in commonly used training dataset non-DBPs at 40% similarity threshold. Finally, non-DBPs with over 40% similarity to at least one DBP of BTD positive set were removed. This step addressed the ambiguity in protein databases, where non-DBPs were not explicitly labeled. By filtering out potential non-DBP sequences with high similarity to known DBPs, we ensure our non-DBP set accurately represents negative examples, minimizing mislabeling. Though it may limit predictions for functionally different yet similar sequences, this trade-off is necessary to ensure reliable benchmarking. This yielded the negative component of BTD comprising 40,734 sequences.

In addition to BTD, we developed two new datasets, EBTD and HBTD to investigate the impact of high and low similarity between training and test datasets on model performance, respectively. While BTD positive and negative sequences have less than 40% similarity compared to the corresponding training sequences, EBTD is the complete opposite in this regard. EBTD creation steps are the same as BTD creation steps except for the boxes in purple as shown in Figure 1. In case of EBTD, we used CD-HIT to retain only those sequences that have over 40% similarity to at least one sequence of the same class from the training set (we made sure that we do not have the same sequence in training and in EBTD). The resulting EBTD contains 2,305 positive samples and 1,963 negative samples. HBTD, on the other hand, is a subset of BTD, with a modification applied during the step marked in purple in Figure 1. Specifically, CD-HIT was used with a 30% similarity threshold instead of the 40% (used for BTD). This adjustment aims to increase the difficulty of predictions, thereby providing a more rigorous evaluation of model robustness. The HBTD dataset consists of 16,003 positive samples and 38,618 negative samples.

Due to emerging needs in the evaluation process, we expanded BTD into two new datasets: BTD-Combo (used in Subsection 6.3) and a proposed train-test dataset (used in Subsection 6.4). BTD-Combo is designed for five-fold cross-validation and comprises non-redundant PDB1075, PDB14K, and BTD. The non-redundancy of PDB1075 and PDB14K was achieved by merging these two datasets and applying CD-HIT with a 40% similarity threshold separately for positive and negative samples (Supplementary Figure S1). This approach ensures minimum data leakage during cross validation using BTD-Combo. BTD-Combo contains 22,332 DBPs and 44,405 non-DBPs. The train-test dataset was constructed from BTD-Combo. We split BTD-Combo into 80% train and 20% test ensuring stratification by class labels and by length percentile groups (Supplementary Figure S1). The training set has 17,857 positive and 35,428 negative sequences while the test set has 4,475 positive and 8,977 negative sequences. This train-test dataset ensures that model performance is evaluated more reliably by minimizing variability due to differences in sequence lengths and class distribution.

## 6. Tool Evaluation

Figure 2 illustrates our tool evaluation pipeline. We start by training the tool models on their respective training datasets from the paper and testing on BTD. The five best tools selected from the this step are then trained on a merged, unbiased training set and tested on BTD, EBTD and HBTD. EBTD testing is performed to show the inflated performance when the training and test datasets are similar, whereas HBTD testing is designed to evaluate the robustness of tools on an even less familiar test dataset. The three best tools selected based on the BTD performance are then five-fold cross-validated on the BTD-Combo dataset (created by merging existing training sets with BTD) using two different class balancing techniques. The goal of this step is to check the influence of training set quality on test performance. The two tools with superior cross-validation performance are then trained and tested based on a length-balanced train-test split created from BTD-Combo. We also benchmark popular bioinformatics tools such as BLAST, ScanProsite, and HMMER on the same train-test split after some simple modifications. Finally, we perform error analysis of the two selected tools based on their test predictions. As evaluation metric, we use sensitivity (true positive rate), specificity (true negative rate) and MCC (Matthews Correlation Coefficient) score (see Supplementary Data 2 for more details of these metrics). We consider DBPs as positive and non-DBPs as negative samples for all cases. The best tools have been chosen based on MCC score in each step of evaluation [72] unless a tool is highly biased towards a particular class (denoted by a large difference between sensitivity and specificity).

**Fig. 2.**
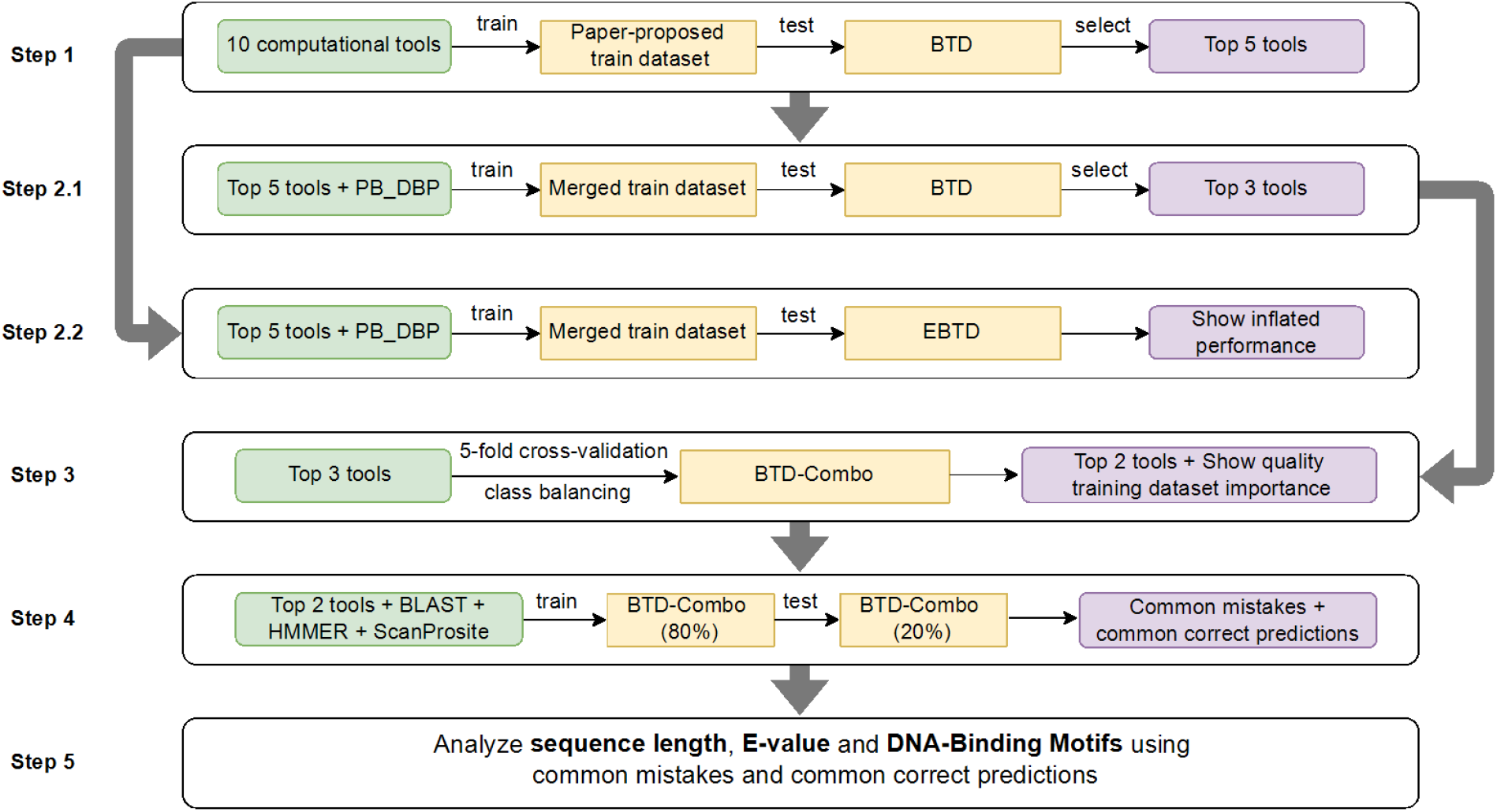
Proposed evaluation pipeline for benchmarking analysis.

### 6.1 Training on Proposed Dataset

We first trained each of the ten tools using the dataset specified in its respective paper. PB_DBP was excluded because its training dataset was not available. This approach ensured that we could replicate the original trained models as closely as possible thereby reproducing the capabilities of each tool. We removed sequences labeled as both positive and negative from these datasets to remove ambiguity. DNABP, Local-DPP, StackDPPred, PDBP-Fusion, LSTM-CNN_Fusion, and PreDBP-PLMs performed the best on the benchmark test set BTD based on MCC score (Table 3). However, the predictions of PreDBP-PLMs were highly biased, exhibiting high specificity but low sensitivity. Therefore, we excluded this tool from further experiments. Conversely, iDNAProt-ES, PseAAC, and DeepDBP showed the worst performance. The possible reasons of their performance issue and potential solutions are discussed in detail in Section 8.

**Table 3.**
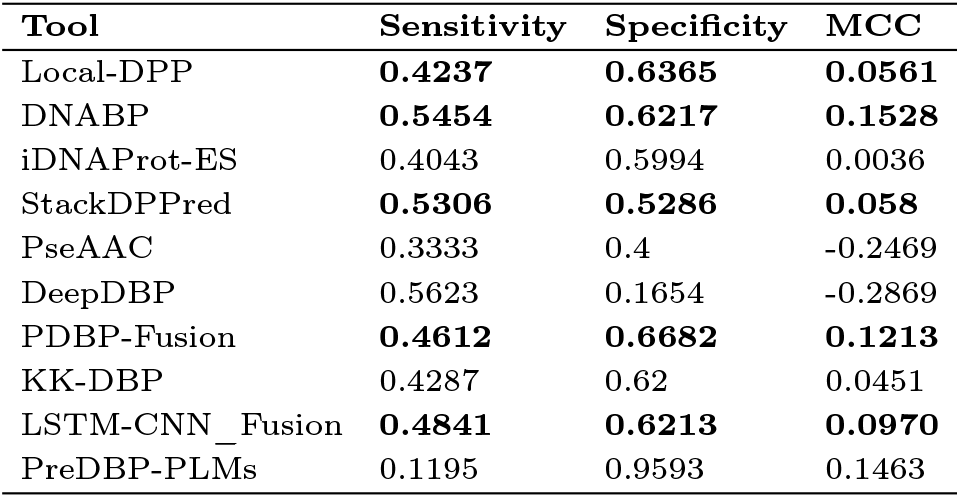
Performance of ten tools on BTD after training on their respective original training data.

### 6.2 Training on Merged Unbiased Dataset

The PDB14K training dataset contains approximately 14 times more samples than PDB1075, which may influence the performance of the corresponding tools due to differences in data distribution and scale. To address this variation in training conditions, we merged PDB14K & PDB1075 and re-evaluated the performance of the top five tools and PB_DBP after training on this new, unified dataset. This approach minimizes discrepancies arising from differences in the size and composition of the original training sets. As shown in Table 4, the average MCC score of the six tools on EBTD is around 0.85 while the score is only around 0.1 on BTD, demonstrating that high similarity between the training and test sets significantly inflates model performance. Such similarity issue is present in the respective original training and test positive samples (DBPs) described in detail in Section 4. We show comparison of the original paper-reported true positive rate (sensitivity) with BTD and EBTD sensitivity in Supplementary Data 5. BTD sensitivity degrades more than 40% on average compared to reported sensitivity, while EBTD sensitivity is within 13% of the reported sensitivity on average. This indicates that the reported performances (Supplementary Data 4) in the original papers are largely inflated and unreliable. We also evaluated the tools on a more challenging test dataset HBTD featuring even less similarity between train and test sequences. The results show that the performance metrics of all six tools dropped slightly. Based on the models’ performance on BTD, we selected the top three tools — Local-DPP, DNABP and LSTM-CNN_Fusion for the next stage of evaluation.

**Table 4.** Performance of six tools trained on the merged unbiased training dataset and tested on BTD, HBTD and EBTD separately.

| Tool | Sensitivity |  |  | Specificity |  |  | MCC |  |  |
| --- | --- | --- | --- | --- | --- | --- | --- | --- | --- |
|  | BT | HT | EBT | BT | HT | EBT | BT | HT | EBT |
| Local-DPP | **0.5168** | 0.5103 | 0.9701 | **0.6642** | 0.6512 | 0.9277 | **0.168** | 0.1503 | 0.9007 |
| DNABP | **0.5422** | 0.537 | 0.9492 | **0.6125** | 0.6044 | 0.8492 | **0.1412** | 0.1297 | 0.8068 |
| StackDPPred | 0.4273 | 0.4187 | 0.9653 | 0.6152 | 0.5972 | 0.9358 | 0.0393 | 0.0148 | 0.9029 |
| PDBP-Fusion | 0.4989 | 0.4932 | 0.9445 | 0.6409 | 0.6316 | 0.8293 | 0.1291 | 0.1158 | 0.7837 |
| LSTM-CNN_Fusion | **0.5023** | 0.4962 | 0.9575 | **0.6491** | 0.6365 | 0.893 | **0.14** | 0.1231 | 0.8553 |
| PB_DBP | 0.4092 | 0.4016 | 0.9562 | 0.7093 | 0.6931 | 0.9032 | 0.1089 | 0.0914 | 0.8631 |

### 6.3 Evaluation after Training Dataset Improvement

Here we explore whether using a more representative training dataset can improve the performance of the top three tools selected in the previous subsection. We implemented five-fold cross-validation on BTD-Combo dataset (described in Section 5). The results in Table 5 indicate a significant improvement in performance for Local-DPP and LSTM-CNN_Fusion, with the new average MCC score exceeding 0.4 suggesting the importance of a more representative training dataset. We also observe the poor performance of DNABP (negative MCC score). DNABP was originally developed based on PDB14K, and certain model parameters can be highly sensitive. With the drastic change in training data, re-tuning of model parameters is often necessary. Furthermore, the features used during this cross-validation were pre-selected based on PDB14K training data used in the corresponding paper, which may cause the performance degradation.

**Table 5.** Five-fold cross-validation performance on original and on balanced BTD-Combo dataset.

| Tool | Imbalanced |  |  | Balanced via Majority Class Undersampling |  |  | Balanced via Minority Class Oversampling |  |  |
| --- | --- | --- | --- | --- | --- | --- | --- | --- | --- |
|  | Sensitivity | Specificity | MCC | Sensitivity | Specificity | MCC | Sensitivity | Specificity | MCC |
| Local-DPP | 0.4541 | 0.9119 | 0.4234 | 0.7148 | 0.7364 | 0.4319 | 0.5283 | 0.8756 | 0.4345 |
| DNABP | 0.1034 | 0.7637 | -0.1591 | 0.3136 | 0.5017 | -0.1757 | 0.1716 | 0.6631 | -0.1733 |
| LSTM-CNN_Fusion | 0.4809 | 0.8766 | 0.3935 | 0.6626 | 0.7659 | 0.4209 | 0.7409 | 0.6956 | 0.415 |

In order to assess the impact of class balancing on model performance, we designed two different class balancing scenarios. We shall discuss their effect on the performance of Local-DPP and LSTM-CNN_Fusion. When training on the original training splits without any balancing, specificity is much higher than sensitivity as the number of non-DBPs is 2X compared to DBPs biasing the model towards negative class prediction. When the majority negative class is randomly undersampled to balance both classes, the sensitivity for both tools increases significantly while decreasing specificity marking a more balanced performance in terms of the two classes. Since a large majority of the protein sequences in the real world are non-DBP (negative class), high true negative rate (specificity) maybe preferred by the scientific community. When the minority positive class is oversampled by repeating each positive sample more than once for class balancing, sensitivity increases in Local-DPP while there is a slight decrease in specificity. This effect seems to be more desirable. Such is not the case for LSTM-CNN_Fusion where there is a large increase in sensitivity and a significant decrease in specificity. Repetitive oversampling makes duplicates of the same samples of the minority class, causing over parameterized deep learning tools such as LSTM-CNN_Fusion to overfit on the minority class, hence such bias is seen towards the minority positive class. Note that throughout the analysis above, the over-and under-sampling were performed only on the training set and not on the test set.

### 6.4. Comparing Against Traditional Tools on Proposed Train-Test Dataset

We now compare Local-DPP and LSTM-CNN_Fusion against traditional tools such as BLAST, ScanProsite and HMMER using our proposed train-test dataset described in Section 5. BLAST and HMMER are both homology detection tools, with BLAST using local alignment-based sequence similarity searching and HMMER employing HMM-based statistical modeling. ScanProsite on the other hand detects homologous regions by finding specific motifs or conserved patterns associated with protein functions. We employed simple repetitive oversampling on the minority positive samples for class balancing during Local-DPP and LSTM-CNN_Fusion model training. Note that we did not perform oversampling on the test set.

To use BLAST for DBP identification, we used the positive subset of the new training dataset as the protein database. Each test sample was queried against this database using BLAST search. If the output list of training DBPs contained a sequence with over 35% identity to the query sequence and an E-value score smaller than a pre-defined threshold (smaller E-value denotes higher similarity) was found for the sequence, the query test sequence was considered positive. Otherwise, it was predicted as negative. Supplementary Data 6 shows BLAST performance in identifying DBPs for different E-value thresholds. We see a decrease in sensitivity and an increase in specificity as we decrease the E-value threshold from 0.01 to 0.000001. We obtained the highest MCC score of 0.402 at a 0.01 threshold (although the MCC scores for the different thresholds are similar). Table 6 shows BLAST performance for the E-value threshold of 0.01. Although the sensitivity of Local-DPP is similar to BLAST, its specificity is better than BLAST. The MCC score of BLAST is slightly smaller compared to both Local-DPP and LSTM-CNN_Fusion. In order to further investigate the perfomance of BLAST, we performed five-fold cross validation on BTD-Combo dataset proposed in Subsection 6.3. We saw similar performance for each of the five folds (Supplementary Data 7) which ensures the consistency of BLAST performance.

**Table 6.** Comparing the best computational tools with traditional tools on proposed train-test dataset.

| Tool | Sensitivity | Specificity | MCC |
| --- | --- | --- | --- |
| Local-DPP | 0.5846 | 0.8501 | 0.4493 |
| LSTM-CNN_Fusion | 0.6974 | 0.7456 | 0.4257 |
| BLAST | 0.5824 | 0.8147 | 0.4020 |
| HMMER | 0.8346 | 0.6038 | 0.4146 |
| ScanProsite | 0.5806 | 0.7918 | 0.3723 |

Our use of HMMER for DBP identification is highly similar to that of BLAST. Specifically, we employed jackhmmer, an iterative search tool that identifies sequence similarities against a protein database. The output of jackhmmer provides hits for each query sequence based on homologous matches in the database. In our approach, we used the positive subset of the train set as the database, and all test sequences were queried against it. If a query sequence had no matches, it was marked as non-DBP. For matched sequences, the lowest E-value of the matching pair was obtained. If the E-value was below the predefined threshold, the sequence was classified as positive (DBP); otherwise, it was considered negative. Supplementary Data 9 shows jackhmmer performance in identifying DBPs for different E-value thresholds. We obtained the highest MCC score of 0.4146 at 0.0001 threshold shown in Table 6. Although HMMER’s MCC score is slightly higher than that of BLAST, its specificity is much lower. Since there are far more negative samples compared to positive ones in real world, we prioritize a tool’s ability to accurately identify non-DBPs. Therefore, we chose to proceed with BLAST for further experiments.

To use ScanProsite for DBP prediction, we simply provided the test sequences to its web server using default settings. ScanProsite compares the submitted sequences against the PROSITE database which utilizes motifs from protein sequences found in UniProtKB, PDB and some other protein databases. For each test sequence, we obtained the motif significance scores for possible DNA-binding motifs identified by ScanProsite. A higher significance score suggests a greater likelihood that the match represents a DNA-binding motif. If more than one such motif was detected, we would assign the highest motif score to the test sequence. In cases where there was no detectable motifs, we simply assigned 0. This assigned score was used for classification based on a pre-defined threshold. ScanProsite achieved the highest MCC score of 0.3723 on test data when we classified a protein as DBP whenever we obtained a positive motif score (last row of Table 6), meaning we were using a classification threshold of 0. ScanProsite’s motif database includes sequences from the Swiss-Prot database, which is also the source of our test data. Hence, there is obvious data leakage from the training to the test dataset. Nevertheless, the two computational tools outperform ScanProsite in terms of MCC score. As we increase the classification threshold to higher values for ScanProsite, the sensitivity keeps going down, while the specificity keeps going up significantly (see Supplementary Data 3).

### 6.5. Combining BLAST with the Best Tools

The top three rows of Table 6 show the best methods according to our evaluation. Let us now consider the prediction of these three methods on our proposed test set described in the previous subsection. Although Local-DPP and LSTM-CNN_Fusion have a considerable amount of overlap in their true positive predictions (%), this overlap is small between these two tools and BLAST (see Venn diagram of Figure 3 and Supplementary Figure S2). Motivated by this discovery, we combined the predictions made by BLAST, Local-DPP and LSTM-CNN_Fusion through majority voting, achieving *sensitivity, specificity and MCC of 0*.*6545, 0*.*8479 and 0*.*5079*, respectively on the proposed test set. This MCC score is significantly higher compared to the best MCC score of 0.4493 achieved by Local-DPP. In fact, this voting system achieves similar high specificity as Local-DPP while achieving a much higher sensitivity. The specificity did not increase, because the overlap of true negative predictions (%) among these three methods is considerably large (Supplementary Figure S3).

**Fig. 3.**
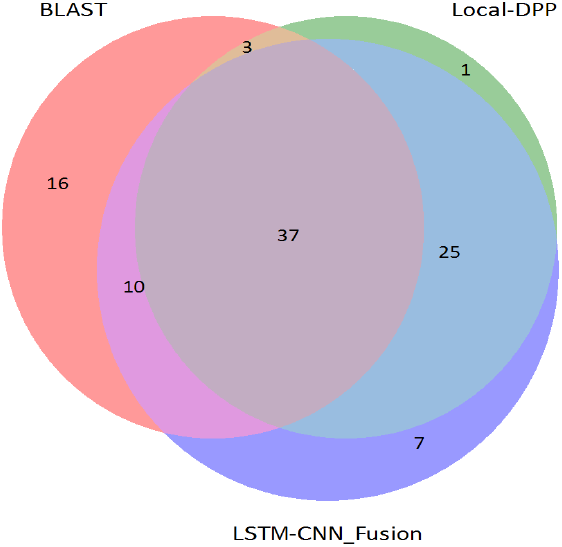
Venn diagram of true positive sequences (%) predicted by BLAST and the two best performing tools.

## 7. Error Analysis

Having identified the best two tools, we proceed to analyze the common mistakes and common correct predictions made by them on the test set described in Supplementary Figure S1. Our aim is to uncover general patterns and challenges inherent in DBP prediction. We discuss the relevant analyses in detail in this section.

### 7.1. Sequence Length Analysis

Figure 4 shows error distributions across different length ranges segmented by every 10^*th*^ percentile of both positive and negative sequences. In case of positive test sequences, shorter sequences exhibit the highest error rates. This error rate consistently decreased with increasing sequence length. Conversely, for negative test sequences, the error rates increased with sequence length. These findings suggest that the selected tools tend to overfit based on the length of training sequences, leading to biased predictions that may obscure the true biological features relevant for classification. The possible reasons of such bias is discussed in the second paragraph of Section 8.

**Fig. 4.**
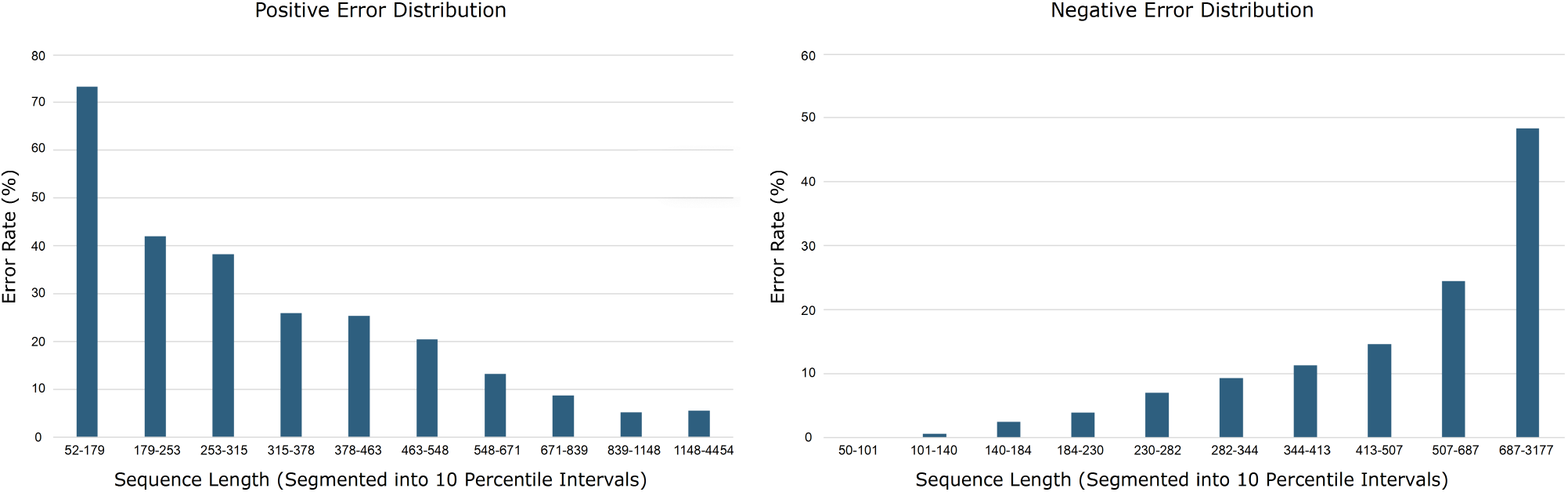
Error rate for positive (left) and negative (right) sequences of different length ranges.

### 7.2. E-ratio Analysis

Before analyzing the E-ratio plots in Figure 5, let us discuss how the E-ratio was generated. The first step was to create two databases named TrainDB+ and TrainDB-containing positive and negative training sequences, respectively. In order to get the E-ratio of a single positive test sample, we performed BLAST search on TrainDB-using the positive test sample as the query and obtained the mean of the lowest 5% E-values (lower E-value indicates higher alignment) denoted as *E-opposite*. We then performed BLAST search on TrainDB+ using the same positive test sample as query sequence and obtained the mean of the lowest 5% E-values denoted as *E-same*. The *E-ratio* of this positive test sample is the ratio of *E-opposite* and *E-same*. This process was repeated for all common correct and common mistake positive test samples to generate the left sub-figure box plots in Figure 5. A higher E-ratio indicates that the positive test sample is more similar to the positive training samples compared to the negative training samples. Similarly, for each negative test sample, we calculated the E-ratio by performing BLAST searches on TrainDB+ to obtain *E-opposite* and on TrainDB-to obtain *E-same*. A higher E-ratio indicates that the negative test sample is more similar to the negative training samples compared to the positive training samples. All E-ratios have been log-scaled for better visualization.

**Fig. 5.**
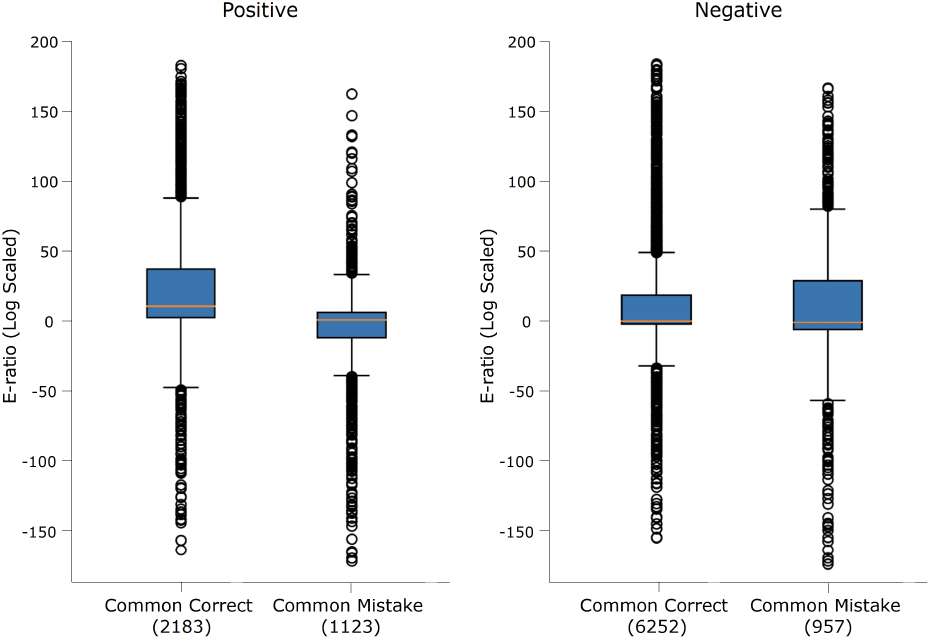
E-ratio plot for positive (left) and negative (right) test samples.

We performed a one-sided Wilcoxon rank-sum test [73], revealing that the E-ratios of correctly classified samples were significantly higher than those of misclassified samples for both the positive and negative classes (p-value of 1.77 *×* 10^*−*85^ and 0.0005 for the positive and negative classes, respectively). This significance analysis indicates that misclassified samples are relatively more similar to training samples of the opposite class compared to the correctly classified samples. This phenomenon is significantly more pronounced in DBPs (positive) compared to non-DBPs (negative).

### 7.3. Motif Score Analysis

Ideally, ScanProsite should not provide any DNA-binding motif as output for non-DBPs; while for DBPs, it should provide one or more motifs with high motif significance scores. Our hypothesis was that mistaken non-DBPs would be assigned relatively high motif scores, while mistaken DBPs would be assigned relatively low motif scores by ScanProsite. Figure 6 shows a summary of the log-scaled maximum motif score for each test sequence (for sequences with no predicted motifs, a log-scaled score of 0 was assigned). Positive (DBP) common correct samples have significantly higher motif scores compared to positive common mistakes (one-sided Wilcoxon rank-sum test p-value of 4.01 *×* 10^*−*140^); while negative common correct samples had much lower motif scores compared to negative common mistakes (one-sided Wilcoxon rank-sum test p-value of 1.54 *×* 10^*−*106^), proving our hypothesis. This means that mistaken non-DBPs have some local patterns which have close resemblance to DNA-binding motifs stored in ScanProsite database, making them harder to predict as non-DBPs.

**Fig. 6.**
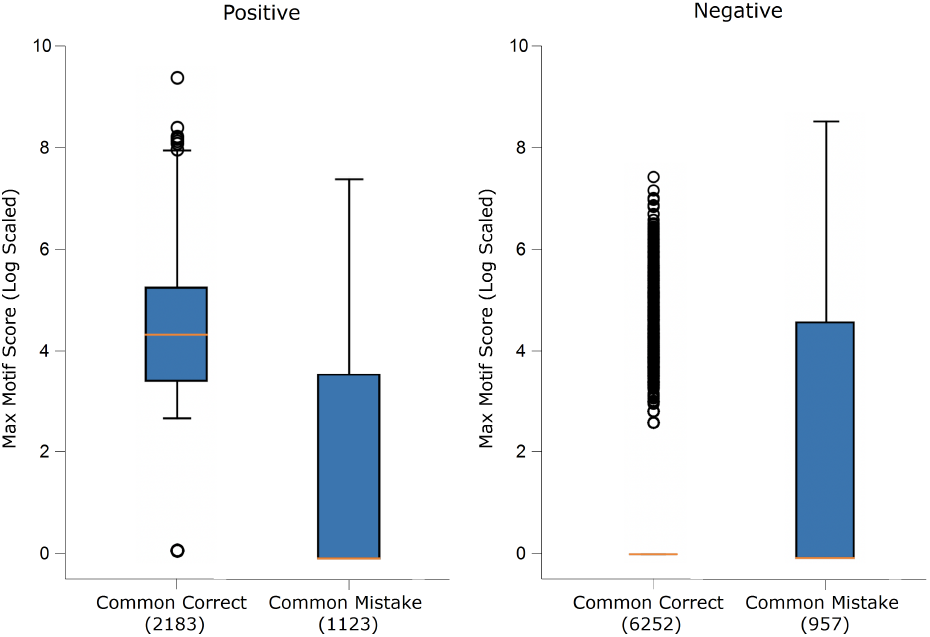
ScanProsite assigned motif score plot for positive (left) and negative (right) test samples.

### 7.4. BLAST-only True Positive Analysis

We examined why 16% of DBPs (652 sequences) were identified only by BLAST and not by the other two tools, as shown in Figure 3. First, we explored why Local-DPP and LSTM-CNN_Fusion failed to correctly classify these sequences by conducting the three error analyses mentioned in the previous three subsections. Plots from the top row of Supplementary Figure S4 indicates that the error pattern of these DBPs for all three analyses are very similar to the commonly mistaken DBPs (positive sequences) by the two tools and hence, they failed to correctly identify these 16% sequences.

Next, to understand why BLAST could identify these DBPs (652 sequences), we investigated how BLAST matches sequences through local alignment. We hypothesized that these sequences were correctly identified due to significant local similarity with known positive sequences from the database. To test this hypothesis, we performed BLAST searches for these 652 sequences (query) against positive training sequences (database), retaining the match with the lowest E-value for each sequence. We then used the Smith-Waterman algorithm [74] to compute the highest local alignment scores for these matches. For comparison, we repeated the same process with two additional sets: the full set of positive sequences correctly and incorrectly predicted by BLAST. Plot from the bottom row of Supplementary Figure S4 shows that the alignment scores of the 652 sequences closely match those of the correctly predicted positives (high alignemnt score) and are distinct from the misclassified ones (low alignemnt score), indicating that local alignment plays a crucial role in BLAST’s predictive success.

## 8. Discussion and Recommendations

Local-DPP and LSTM-CNN_Fusion are the two best tools according to our benchmarking. Both of these tools used evolutionary features generated from PSSM transformation, highlighting the importance of such features. LSTM-CNN_Fusion truncates the PSSM matrix to a fixed size, while Local-DPP calculates summary statistics for a Random Forest model. Such processing may lead to significant loss of information. Instead, these variable-size feature matrices might be more smartly processed utilizing recent variable-size graph learning algorithms [75, 76]. Additionally, instead of using BLAST for PSSM generation, software such as HHblits might be used for obtaining profile HMMs (more generalized form of PSSMs) from sequences [77]. Such measures might help reduce the E-ratio bias described in Subsection 7.2.

For the two recent deep learning-based tools named PDBP-Fusion and LSTM-CNN_Fusion, a notable issue is the need to select a fixed maximum length (700) for input sequences to fit within the neural network’s architecture. This constraint might result in the loss of crucial information, as sequences exceeding the predetermined length are truncated. Furthermore, these tools take a residue-based encoding scheme when using long short term memory (LSTM) model, meaning that the LSTM has to go through 700 time steps before making a prediction. The models were originally trained on only around 14K training samples while needing to learn sequential patterns over a path of length 700. Such a model might cause sub-optimal learning and overfitting [78, 79]. In such cases, smart window-based encoding schemes [80] and attention mechanisms [81, 82] might help in reducing the length bias described in Subsection 7.1. This length bias also exists in random forest based Local-DPP, probably because the size of the sub-PSSMs it uses as its features depend on the protein sequence length.

The latest tools, PB_DBP and PreDBP-PLMs, highlight the trend of performing bioinformatics prediction tasks with the help of PLMs [83, 84, 85, 86]. While promising, they tend to have high specificity and very low sensitivity, leading to the misclassification of many actual DBPs. Vast majority of the sequences used for PLM pretraining consists of non-DBPs. Since these models are not fine-tuned on DBP identification specific training data in PB_DBP and PreDBP-PLMs, they may generate embeddings that fail to carry DBP-specific features. Future work should focus on refining these pretrained PLMs to reduce biases and improve generalization.

DeepDBP, PseAAC and iDNAProt-ES are the three tools showing the worst performance during our benchmarking. Both DeepDBP and PseAAC relied solely on sequence-derived summary statistics, which shows the necessity of more informative novel sequence-derived features and perhaps also inclusion of evolutionary and structural features. While exploring ways to incorporate structural insights, we considered using AlphaFold [87], which tackles a key challenge in molecular biology: predicting protein structures from sequence data, a task traditionally reliant on costly and time-intensive methods like X-ray crystallography or NMR spectroscopy [88]. By leveraging machine learning, AlphaFold offers rapid and accurate predictions, greatly advancing structural biology. Structural insights from AlphaFold can enhance DBP prediction by capturing subtle structural motifs and spatial interactions that are difficult to infer from sequence data alone. However, despite its potential, AlphaFold’s large runtime and computational cost (exceeding one hour per sequence on T4 GPU) may make it challenging to use for large-scale studies and to deploy for real world use. On the other hand, iDNAProt-ES used all three types of features (sequence, evolutionary and structural), but still demonstrated poor performance. This particular method derived many different features from the PSSMs and from the torsion angles of the structural features, and might have suffered from errors in the determination (by third-party tools) of these features and from a severely increased curse of dimensionality. Using irrelevant and/or error-containing features might lead to overfitting [54, 89].

All tools utilizing classic ML models require one single feature vector per protein sequence. As a result, heterogeneous features of different scales and different inter-sample differences are concatenated together leading to inaccurate model behavior [90, 91]. Instead, each heterogeneous feature type might be encoded in different branches of a learnable network as suggested in [92]. Tools such as StackDPPred stack various ML models in multiple stages, which may introduce additional complexity without substantial benefit. Proper ablation studies should be conducted before implementing stacking strategies to ensure they are justified [93, 94]. There are tools such as HMMER and ScanProsite that provide potential DNA-binding motif locations along with significance scores in the protein sequences. These locations might be given special importance during DBP prediction. Furthermore, tools like DNABR [48], DRNApred [95] and HybridDBRpred [96] might detect whether a residue of a protein is DNA-binding or not. Such residue-level predictions might assist in modeling the overall behavior of the entire protein sequence, thereby reducing the motif significance bias described in Subsection 7.3.

Our evaluation showed moderate performance gain of the two best computational tools over traditional BLAST in terms of MCC. However, many DBPs predicted by them were not predicted by BLAST. The ability to predict DBPs missed by BLAST adds value to these new tools. Indeed, combining these three tools through simple means such as majority voting achieved significantly better performance (Subsection 6.5). This simple ensemble approach achieved 65% sensitivity and 85% specificity on our proposed test set. From the literature [22], it can be estimated that there are 9X more non-DBPs (negative class) compared to DBPs (positive class) in real-life. Thus, every DBP correctly identified by this ensemble approach would be accompanied by about two false positives. This might be good enough for practical use, though there is scope for further improvement.

Researchers should avoid data leakage between training and test set through proper similarity threshold based filtering. Test set should be representative of the real world in terms of sample number, heterogeneity and class imbalance. We have provided our developed training and test fasta files via GitHub. Researchers can perform cross-validation on the training set to develop their models, while the test set can be used for final validation. Note that both the training and test data we have provided contain significantly more non-DBPs compared to DBPs. All the recent tools we benchmarked have recommended using random undersampling for class balancing the training set. However, as shown in Table 5, this balancing approach negatively impacts the true negative rate (specificity). Random undersampling of the majority class can potentially introduce gaps in the feature space making the trained model less robust in such areas [97, 98]. Instead, clustering-based undersampling might be performed to ensure that sampling covers the entire majority class feature space [99]. Alternatively, oversampling of the minority class might be preferable.

One prominent issue is the lack of usability of these developed tools. Although some of the tools provide their own web server, these servers are rarely maintained after publication; as they are not commercial software. The best practice would be to provide the codes along with the trained model via free public repositories such as GitHub, GitLab or Zenodo such that it is possible to run these tools with minimal effort on raw sequences. We have provided Local-DPP, LSTM-CNN_Fusion, BLAST and majority voting based ensembling scheme for DBP identification via GitHub which can be followed as an example by future researchers.

In our benchmarking approach, we show that many existing works on DBP identification have not been thoughtfully tested. A more thoughtfully designed evaluation is then presented and these existing works are then tested accordingly, revealing that their previously reported performance is exaggerated due to the effects of data doppelgangers in their test sets and using insufficiently representative training and test datasets. By retraining on a more representative training dataset, two of these previously reported methods are “rescued” in terms of performance, though they did not perform significantly better than BLAST. The true value of the retrained methods is then demonstrated by showing that their predicted sets of DBPs are distinct from those identified by BLAST; and thus, a simple majority vote among these two retrained methods and BLAST yields superior performance. Poor evaluation design can be observed in many other protein class prediction problems, and similarly, many methods proposed for these problems may have reported seemingly exaggerated performance (likely for similar reasons of data doppelgangers and non-representative training/test datasets). The same benchmarking strategy should be readily applicable to studying these problems. Examples of such problems include (but are not restricted to) -(a) protein function prediction [100], (b) subcellular localization prediction [101], (c) protein-protein interaction prediction [102], and (d) post translational modification site prediction [103].

## 9. Conclusion

We benchmarked eleven state-of-the-art computational DBP identification tools in this paper. We began by categorizing and analyzing these tools based on their models, datasets and types of features used. By scrutinizing the conventional datasets commonly used by these tools, we identified significant limitations, particularly issues related to data leakage leading to inflated performance. To address these issues, we developed two new benchmarking datasets: BTD and EBTD. We demonstrated the inflated performance using EBTD. BTD, designed to mitigate the adverse effects of data leakage, provides a more realistic and unbiased evaluation of different DBP prediction tools, serving as a valuable reference for future tool development and evaluation. Using BTD, we re-evaluated the eleven tools, selected the best two tools, and assessed their effectiveness against traditional methods in predicting DBPs through simple adjustments. We showed the significant performance gain of the two tools combined with traditional BLAST search. Additionally, we provided a high-quality train-test dataset for future development based on BTD and available popular training datasets. This dataset along with the top-performing methods (Local-DPP, LSTM-CNN_Fusion, BLAST) and their ensemble classifier are publicly available at https://github.com/Rafeed-bot/DNA_BP_Benchmarking. These methods are directly applicable on raw protein sequences for DBP identification. Beyond tool evaluation, we analyzed the mistakes made by top-performing tools, providing insights for improvement of DBP prediction tools and explored the reasons why BLAST outperformed these tools on certain positive samples. Finally, we discussed possible limitations of current models, feature extraction methods and data balancing techniques, and offered potential solutions for future research efforts in this field.

## References

1. Christoph Zimmer and Ulla Wähnert. Nonintercalating dna-binding ligands: specificity of the interaction and their use as tools in biophysical, biochemical and biological investigations of the genetic material. Progress in biophysics and molecular biology, 47(1):31–112, 1986.

2. R G Brennan and B W Matthews. The helix-turn-helix dna binding motif. Journal of Biological Chemistry, 264(4):1903–1906, 1989.

3. Robert A Moxley, Harry W Jarrett, and Suchareeta Mitra. Methods for transcription factor separation. Journal of Chromatography B, 797(1):269–288, 2003. Interactions in Biological Systems.

4. Aaron Klug. The discovery of zinc fingers and their applications in gene regulation and genome manipulation. Annual review of biochemistry, 79(1):213–231, 2010.

5. David S Latchman. Transcription factors: an overview. The international journal of biochemistry & cell biology, 29(12):1305–1312, 1997.

6. Karolin Luger, Armin W Mäder, Robin K Richmond, David F Sargent, and Timothy J Richmond. Crystal structure of the nucleosome core particle at 2.8 å resolution. Nature, 389(6648):251–260, 1997.

7. Stefan Oehler, Regina Alex, and Andrew Barker. Is nitrocellulose filter binding really a universal assay for protein–dna interactions? Analytical biochemistry, 268(2):330–336, 1999.

8. Katie Freeman, Marc Gwadz, and David Shore. Molecular and genetic analysis of the toxic effect of rap1 overexpression in yeast. Genetics, 141(4):1253–1262, 1995.

9. Michael J Buck and Jason D Lieb. Chip-chip: considerations for the design, analysis, and application of genome-wide chromatin immunoprecipitation experiments. Genomics, 83(3):349–360, 2004.

10. Robert E Langlois and Hui Lu. Boosting the prediction and understanding of dna-binding domains from sequence. Nucleic acids research, 38(10):3149–3158, 2010.

11. Jay Shendure and Hanlee Ji. Next-generation dna sequencing. Nature biotechnology, 26(10):1135–1145, 2008.

12. Yu-dong Cai and Shuo Liang Lin. Support vector machines for predicting rrna-, rna-, and dna-binding proteins from amino acid sequence. Biochimica et Biophysica Acta (BBA)-Proteins and Proteomics, 1648(1-2):127–133, 2003.

13. Nitin Bhardwaj, Robert E Langlois, Guijun Zhao, and Hui Lu. Kernel-based machine learning protocol for predicting dna-binding proteins. Nucleic Acids Research, 33(20):6486–6493, 2005.

14. Gajendra PS Raghava, Michael M Gromiha, and Manish Kumar. Identification of dna-binding proteins using support vector machines and evolutionary profiles. 2007.

15. Hui-Lin Huang, I-Che Lin, Yi-Fan Liou, Chia-Ta Tsai, Kai-Ti Hsu, Wen-Lin Huang, Shinn-Jang Ho, and Shinn-Ying Ho. Predicting and analyzing dna-binding domains using a systematic approach to identifying a set of informative physicochemical and biochemical properties. Bmc Bioinformatics, 12:1–13, 2011.

16. Yanping Zhang, Jun Xu, Wei Zheng, Chen Zhang, Xingye Qiu, Ke Chen, and Jishou Ruan. newdna-prot: Prediction of dna-binding proteins by employing support vector machine and a comprehensive sequence representation. Computational biology and chemistry, 52:51–59, 2014.

17. Jiansheng Wu, Hongde Liu, Xueye Duan, Yan Ding, Hongtao Wu, Yunfei Bai, and Xiao Sun. Prediction of dna-binding residues in proteins from amino acid sequences using a random forest model with a hybrid feature. Bioinformatics, 25(1):30–35, 2009.

18. Guy Nimrod, Maya Schushan, András Szilágyi, Christina Leslie, and Nir Ben-Tal. idbps: a web server for the identification of dna binding proteins. Bioinformatics, 26(5):692–693, 2010.

19. M Saifur Rahman, Swakkhar Shatabda, Sanjay Saha, Mohammad Kaykobad, and M Sohel Rahman. Dpp-pseaac: a dna-binding protein prediction model using chou’s general pseaac. Journal of theoretical biology, 452:22–34, 2018.

20. Ziliang Qian, Yu-Dong Cai, and Yixue Li. A novel computational method to predict transcription factor dna binding preference. Biochemical and biophysical research communications, 348(3):1034–1037, 2006.

21. Babak Alipanahi, Andrew Delong, Matthew T Weirauch, and Brendan J Frey. Predicting the sequence specificities of dna-and rna-binding proteins by deep learning. Nature biotechnology, 33(8):831–838, 2015.

22. Yu-Hui Qu, Hua Yu, Xiu-Jun Gong, Jia-Hui Xu, and Hong-Shun Lee. On the prediction of dna-binding proteins only from primary sequences: A deep learning approach. PloS one, 12(12):e0188129, 2017.

23. Weizhong Lu, Xiaoyi Chen, Yu Zhang, Hongjie Wu, Yijie Ding, Jiawei Shen, Shixuan Guan, and Haiou Li. Research on dna-binding protein identification method based on lstm-cnn feature fusion. Computational and Mathematical Methods in Medicine, 2022(1):9705275, 2022.

24. Guobin Li, Xiuquan Du, Xinlu Li, L. Zou, Guanhong Zhang, and Zhize Wu. Prediction of dna binding proteins using local features and long-term dependencies with primary sequences based on deep learning. PeerJ, 9:e11262, 2021.

25. Ahmed Elnaggar, Michael Heinzinger, Christian Dallago, Ghalia Rehawi, Yu Wang, Llion Jones, Tom Gibbs, Tamas Feher, Christoph Angerer, Martin Steinegger, et al. Prottrans: Toward understanding the language of life through self-supervised learning. IEEE transactions on pattern analysis and machine intelligence, 44(10):7112–7127, 2021.

26. Roshan Rao, Nicholas Bhattacharya, Neil Thomas, Yan Duan, Peter Chen, John Canny, Pieter Abbeel, and Yun Song. Evaluating protein transfer learning with tape. Advances in neural information processing systems, 32, 2019.

27. A Vaswani. Attention is all you need. Advances in Neural Information Processing Systems, 2017.

28. András Szilágyi and Jeffrey Skolnick. Efficient prediction of nucleic acid binding function from low-resolution protein structures. Journal of molecular biology, 358(3):922–933, 2006.

29. K Krishna Kumar, Ganesan Pugalenthi, and Ponnuthurai N Suganthan. Dna-prot: identification of dna binding proteins from protein sequence information using random forest. Journal of Biomolecular Structure and Dynamics, 26(6):679–686, 2009.

30. Y Fang, Y Guo, Y Feng, and M Li. Predicting dna-binding proteins: approached from chou’s pseudo amino acid composition and other specific sequence features. Amino acids, 34:103–109, 2008.

31. Loris Nanni and Alessandra Lumini. Combing ontologies and dipeptide composition for predicting dna-binding proteins. Amino Acids, 34:635–641, 2008.

32. Li Song, Dapeng Li, Xiangxiang Zeng, Yunfeng Wu, Li Guo, and Quan Zou. ndna-prot: identification of dna-binding proteins based on unbalanced classification. BMC bioinformatics, 15:1–10, 2014.

33. Bin Liu, Jinghao Xu, Shixi Fan, Ruifeng Xu, Jiyun Zhou, and Xiaolong Wang. Psedna-pro: Dna-binding protein identification by combining chou’s pseaac and physicochemical distance transformation. Molecular Informatics, 34(1):8–17, 2015.

34. Dawei Qi, Chen Song, and Taigang Liu. Predbp-plms: Prediction of dna-binding proteins based on pre-trained protein language models and convolutional neural networks. Analytical Biochemistry, 694:115603, 2024.

35. Jinfeng Li, Shun Zhang, and Chun Fang. Pb_dbp: Identifying dna-binding proteins using probert_bilstm model. In Proceedings of the 2023 6th International Conference on Big Data Technologies, pages 242–246, 2023.

36. Wangchao Lou, Xiaoqing Wang, Fan Chen, Yixiao Chen, Bo Jiang, and Hua Zhang. Sequence based prediction of dna-binding proteins based on hybrid feature selection using random forest and gaussian naive bayes. PloS one, 9(1):e86703, 2014.

37. Shahana Yasmin Chowdhury, Swakkhar Shatabda, and Abdollah Dehzangi. idnaprot-es: identification of dna-binding proteins using evolutionary and structural features. Scientific reports, 7(1):14938, 2017.

38. Farman Ali, Saeed Ahmed, Zar Nawab Khan Swati, and Shahid Akbar. Dp-binder: machine learning model for prediction of dna-binding proteins by fusing evolutionary and physicochemical information. Journal of Computer-Aided Molecular Design, 33:645–658, 2019.

39. Stephen F Altschul, Warren Gish, Webb Miller, Eugene W Myers, and David J Lipman. Basic local alignment search tool. Journal of molecular biology, 215(3):403–410, 1990.

40. Edouard De Castro, Christian JA Sigrist, Alexandre Gattiker, Virginie Bulliard, Petra S Langendijk-Genevaux, Elisabeth Gasteiger, Amos Bairoch, and Nicolas Hulo. Scanprosite: detection of prosite signature matches and prorule-associated functional and structural residues in proteins. Nucleic acids research, 34(suppl_2):W362–W365, 2006.

41. Sean R Eddy. Accelerated profile hmm searches. PLoS computational biology, 7(10):e1002195, 2011.

42. Leyi Wei, Jijun Tang, and Quan Zou. Local-dpp: An improved dna-binding protein prediction method by exploring local evolutionary information. Information Sciences, 384:135–144, 2017.

43. Xin Ma, Jing Guo, and Xiao Sun. Dnabp: Identification of dna-binding proteins based on feature selection using a random forest and predicting binding residues. PloS one, 11(12):e0167345, 2016.

44. Avdesh Mishra, Pujan Pokhrel, and Md Tamjidul Hoque. Stackdppred: a stacking based prediction of dna-binding protein from sequence. Bioinformatics, 35(3):433–441, 2019.

45. Sheikh Adilina, Dewan Md Farid, and Swakkhar Shatabda. Effective dna binding protein prediction by using key features via chou’s general pseaac. Journal of theoretical biology, 460:64–78, 2019.

46. Shadman Shadab, Md Tawab Alam Khan, Nazia Afrin Neezi, Sheikh Adilina, and Swakkhar Shatabda. Deepdbp: deep neural networks for identification of dna-binding proteins. Informatics in Medicine Unlocked, 19:100318, 2020.

47. Yuran Jia, Shan Huang, and Tianjiao Zhang. Kk-dbp: a multi-feature fusion method for dna-binding protein identification based on random forest. Frontiers in Genetics, 12:811158, 2021.

48. Xin Ma, Jing Guo, Hong-De Liu, Jian-Ming Xie, and Xiao Sun. Sequence-based prediction of dna-binding residues in proteins with conservation and correlation information. IEEE/ACM transactions on computational biology and bioinformatics, 9(06):1766–1775, 2012.

49. Christian B Anfinsen. Principles that govern the folding of protein chains. Science, 181(4096):223–230, 1973.

50. Ken A Dill and Justin L MacCallum. The protein-folding problem, 50 years on. science, 338(6110):1042–1046, 2012.

51. Stephen F Altschul, Thomas L Madden, Alejandro A Schäffer, Jinghui Zhang, Zheng Zhang, Webb Miller, and David J Lipman. Gapped blast and psi-blast: a new generation of protein database search programs. Nucleic acids research, 25(17):3389–3402, 1997.

52. Yuedong Yang, Rhys Heffernan, Kuldip Paliwal, James Lyons, Abdollah Dehzangi, Alok Sharma, Jihua Wang, Abdul Sattar, and Yaoqi Zhou. Spider2: a package to predict secondary structure, accessible surface area, and main-chain torsional angles by deep neural networks. Prediction of protein secondary structure, pages 55–63, 2017.

53. Zsuzsanna Dosztanyi, Veronika Csizmok, Peter Tompa, and Istvan Simon. The pairwise energy content estimated from amino acid composition discriminates between folded and intrinsically unstructured proteins. Journal of molecular biology, 347(4):827–839, 2005.

54. Isabelle Guyon and André Elisseeff. An introduction to variable and feature selection. Journal of machine learning research, 3(Mar):1157–1182, 2003.

55. Leo Breiman. Random forests. Machine learning, 45:5–32, 2001.

56. Corinna Cortes and Vladimir Vapnik. Support-vector networks. Machine learning, 20:273–297, 1995.

57. Trevor Hastie, Robert Tibshirani, Jerome H Friedman, and Jerome H Friedman. The elements of statistical learning: data mining, inference, and prediction, volume 2. Springer, 2009.

58. Naomi S Altman. An introduction to kernel and nearest-neighbor nonparametric regression. The American Statistician, 46(3):175–185, 1992.

59. Bernhard Schölkopf and Alexander J Smola. Learning with kernels: support vector machines, regularization, optimization, and beyond. MIT press, 2002.

60. Yann LeCun, Léon Bottou, Yoshua Bengio, and Patrick Haffner. Gradient-based learning applied to document recognition. Proceedings of the IEEE, 86(11):2278–2324, 1998.

61. Sepp Hochreiter and Jürgen Schmidhuber. Long short-term memory. Neural computation, 9(8):1735–1780, 1997.

62. Bin Liu, Jinghao Xu, Xun Lan, Ruifeng Xu, Jiyun Zhou, Xiaolong Wang, and Kuo-Chen Chou. idna-prot| dis: identifying dna-binding proteins by incorporating amino acid distance-pairs and reduced alphabet profile into the general pseudo amino acid composition. PloS one, 9(9):e106691, 2014.

63. Helen M Berman, John Westbrook, Zukang Feng, Gary Gilliland, Talapady N Bhat, Helge Weissig, Ilya N Shindyalov, and Philip E Bourne. The protein data bank. Nucleic acids research, 28(1):235–242, 2000.

64. Guoli Wang and Roland L Dunbrack. Pisces: recent improvements to a pdb sequence culling server. Nucleic acids research, 33(suppl_2):W94–W98, 2005.

65. Nathalie Japkowicz. The class imbalance problem: Significance and strategies. In Proc. of the Int’l Conf. on artificial intelligence, volume 56, pages 111–117, 2000.

66. Haibo He and Edwardo A Garcia. Learning from imbalanced data. IEEE Transactions on knowledge and data engineering, 21(9):1263–1284, 2009.

67. Weizhong Li and Adam Godzik. Cd-hit: a fast program for clustering and comparing large sets of protein or nucleotide sequences. Bioinformatics, 22(13):1658–1659, 2006.

68. Li Rong Wang, Limsoon Wong, and Wilson Wen Bin Goh. How doppelgänger effects in biomedical data confound machine learning. Drug discovery today, 27(3):678–685, 2022.

69. Mengyuan Yang, Huifen Lu, Jiajia Liu, Sijia Wu, Pora Kim, and Xiaobo Zhou. lncrnafunc: a knowledgebase of lncrna function in human cancer. Nucleic acids research, 50(D1):D1295–D1306, 2022.

70. Jianzhi Zhang. Evolution by gene duplication: an update. Trends in ecology & evolution, 18(6):292–298, 2003.

71. Uniprot: the universal protein knowledgebase in 2023. Nucleic acids research, 51(D1):D523–D531, 2023.

72. Davide Chicco and Giuseppe Jurman. The advantages of the matthews correlation coefficient (mcc) over f1 score and accuracy in binary classification evaluation. BMC genomics, 21:1–13, 2020.

73. Frank Wilcoxon. Individual comparisons by ranking methods. In Breakthroughs in statistics: Methodology and distribution, pages 196–202. Springer, 1992.

74. Temple F Smith, Michael S Waterman, et al. Identification of common molecular subsequences. Journal of molecular biology, 147(1):195–197, 1981.

75. Thomas N Kipf and Max Welling. Semi-supervised classification with graph convolutional networks. arXiv preprint arXiv:1609.02907, 2016.

76. Petar Velickovic, Guillem Cucurull, Arantxa Casanova, Adriana Romero, Pietro Lio, Yoshua Bengio, et al. Graph attention networks. stat, 1050(20):10–48550, 2017.

77. Michael Remmert, Andreas Biegert, Andreas Hauser, and Johannes Söding. Hhblits: lightning-fast iterative protein sequence searching by hmm-hmm alignment. Nature methods, 9(2):173–175, 2012.

78. Razvan Pascanu, Tomas Mikolov, and Yoshua Bengio. On the difficulty of training recurrent neural networks. In International conference on machine learning, pages 1310–1318. Pmlr, 2013.

79. Klaus Greff, Rupesh K Srivastava, Jan Koutník, Bas R Steunebrink, and Jürgen Schmidhuber. Lstm: A search space odyssey. IEEE transactions on neural networks and learning systems, 28(10):2222–2232, 2016.

80. Jessica Lin, Eamonn Keogh, Stefano Lonardi, and Bill Chiu. A symbolic representation of time series, with implications for streaming algorithms. In Proceedings of the 8th ACM SIGMOD workshop on Research issues in data mining and knowledge discovery, pages 2–11, 2003.

81. Dzmitry Bahdanau, Kyunghyun Cho, and Yoshua Bengio. Neural machine translation by jointly learning to align and translate. arXiv preprint arXiv:1409.0473, 2014.

82. Minh-Thang Luong, Hieu Pham, and Christopher D Manning. Effective approaches to attention-based neural machine translation. arXiv preprint arXiv:1508.04025, 2015.

83. Amelia Villegas-Morcillo, Angel M Gomez, and Victoria Sanchez. An analysis of protein language model embeddings for fold prediction. Briefings in Bioinformatics, 23(3):bbac142, 2022.

84. Konstantin Weissenow, Michael Heinzinger, and Burkhard Rost. Protein language-model embeddings for fast, accurate, and alignment-free protein structure prediction. Structure, 30(8):1169–1177, 2022.

85. Yue Zhang, Jianyuan Lin, Lianmin Zhao, Xiangxiang Zeng, and Xiangrong Liu. A novel antibacterial peptide recognition algorithm based on bert. Briefings in bioinformatics, 22(6):bbab200, 2021.

86. Qianmu Yuan, Sheng Chen, Yu Wang, Huiying Zhao, and Yuedong Yang. Alignment-free metal ion-binding site prediction from protein sequence through pretrained language model and multi-task learning. Briefings in bioinformatics, 23(6):bbac444, 2022.

87. John Jumper, Richard Evans, Alexander Pritzel, Tim Green, Michael Figurnov, Olaf Ronneberger, Kathryn Tunyasuvunakool, Russ Bates, Augustin Žídek, Anna Potapenko, et al. Highly accurate protein structure prediction with alphafold. nature, 596(7873):583–589, 2021.

88. Letícia MF Bertoline, Angélica N Lima, Jose E Krieger, and Samantha K Teixeira. Before and after alphafold2: An overview of protein structure prediction. Frontiers in bioinformatics, 3:1120370, 2023.

89. Trevor Hastie. The elements of statistical learning: data mining, inference, and prediction, 2009.

90. Jair Cervantes, Farid Garcia-Lamont, Lisbeth Rodríguez-Mazahua, and Asdrubal Lopez. A comprehensive survey on support vector machine classification: Applications, challenges and trends. Neurocomputing, 408:189–215, 2020.

91. Gilles Louppe, Louis Wehenkel, Antonio Sutera, and Pierre Geurts. Understanding variable importances in forests of randomized trees. Advances in neural information processing systems, 26, 2013.

92. Chowdhury Rafeed Rahman, Ruhul Amin, Swakkhar Shatabda, and Md Sadrul Islam Toaha. A convolution based computational approach towards dna n6-methyladenine site identification and motif extraction in rice genome. Scientific Reports, 11(1):10357, 2021.

93. Mark J Van der Laan, Eric C Polley, and Alan E Hubbard. Super learner. Statistical applications in genetics and molecular biology, 6(1), 2007.

94. Joseph Sill, Gábor Takács, Lester Mackey, and David Lin. Feature-weighted linear stacking. arXiv preprint arXiv:0911.0460, 2009.

95. Jing Yan and Lukasz Kurgan. Drnapred, fast sequence-based method that accurately predicts and discriminates dna-and rna-binding residues. Nucleic acids research, 45(10):e84–e84, 2017.

96. Jian Zhang, Sushmita Basu, and Lukasz Kurgan. Hybriddbrpred: improved sequence-based prediction of dna-binding amino acids using annotations from structured complexes and disordered proteins. Nucleic Acids Research, 52(2):e10–e10, 2024.

97. Gustavo EAPA Batista, Ronaldo C Prati, and Maria Carolina Monard. A study of the behavior of several methods for balancing machine learning training data. ACM SIGKDD explorations newsletter, 6(1):20–29, 2004.

98. Haibo He and Edwardo A Garcia. Learning from imbalanced data. IEEE Transactions on knowledge and data engineering, 21(9):1263–1284, 2009.

99. Show-Jane Yen and Yue-Shi Lee. Cluster-based under-sampling approaches for imbalanced data distributions. Expert Systems with Applications, 36(3):5718–5727, 2009.

100. Maxat Kulmanov and Robert Hoehndorf. Deepgoplus: improved protein function prediction from sequence. Bioinformatics, 36(2):422–429, 2020.

101. José Juan Almagro Armenteros, Casper Kaae Sønderby, Søren Kaae Sønderby, Henrik Nielsen, and Ole Winther. Deeploc: prediction of protein subcellular localization using deep learning. Bioinformatics, 33(21):3387–3395, 2017.

102. Xiuquan Du, Shiwei Sun, Changlin Hu, Yu Yao, Yuanting Yan, and Yanping Zhang. Deepppi: boosting prediction of protein–protein interactions with deep neural networks. Journal of chemical information and modeling, 57(6):1499–1510, 2017.

103. Fenglin Luo, Minghui Wang, Yu Liu, Xing-Ming Zhao, and Ao Li. Deepphos: prediction of protein phosphorylation sites with deep learning. Bioinformatics, 35(16):2766–2773, 2019.

