## Supplementary Figures for "Benchmarking Recent Computational Tools for DNA-binding Protein Identification"

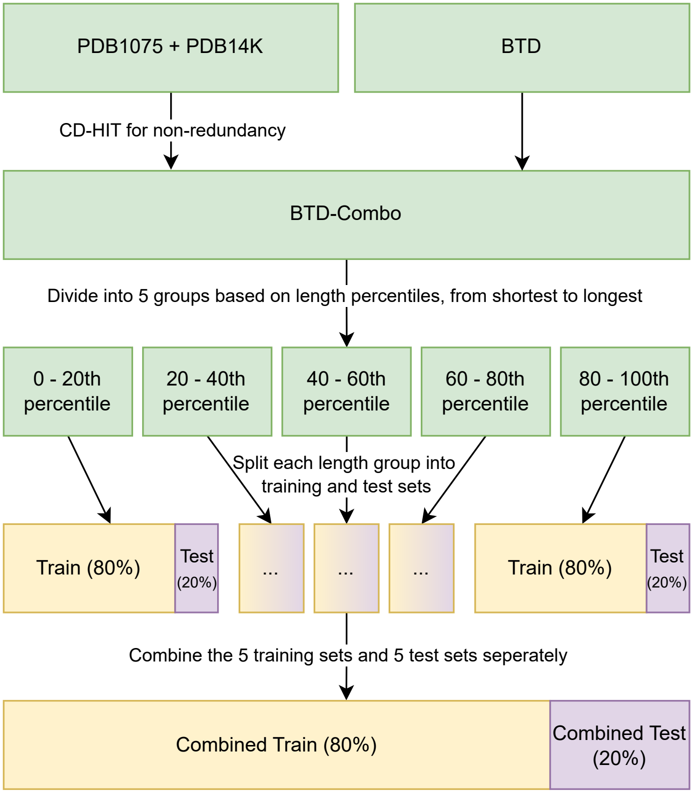


Fig S1: Proposed Train-Test Split Creation Process

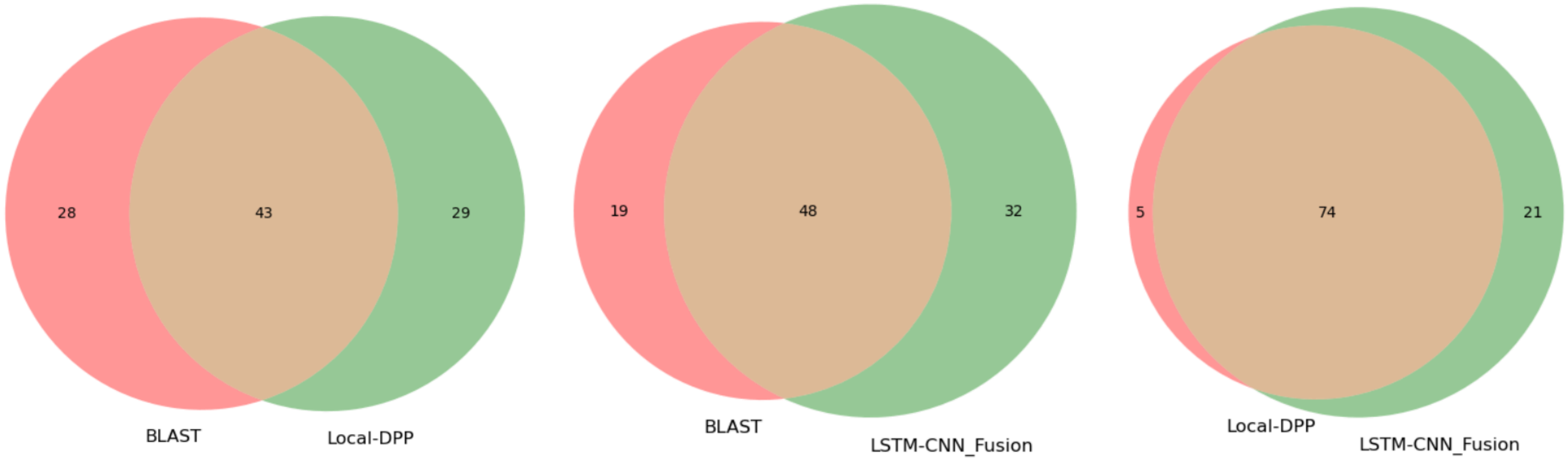


*Fig S2: Venn diagram of true positive sequences for all combinations of tool pairs for the three tools*

*Fig S3: Venn diagrams for the true negative sequences predicted by the three tools*

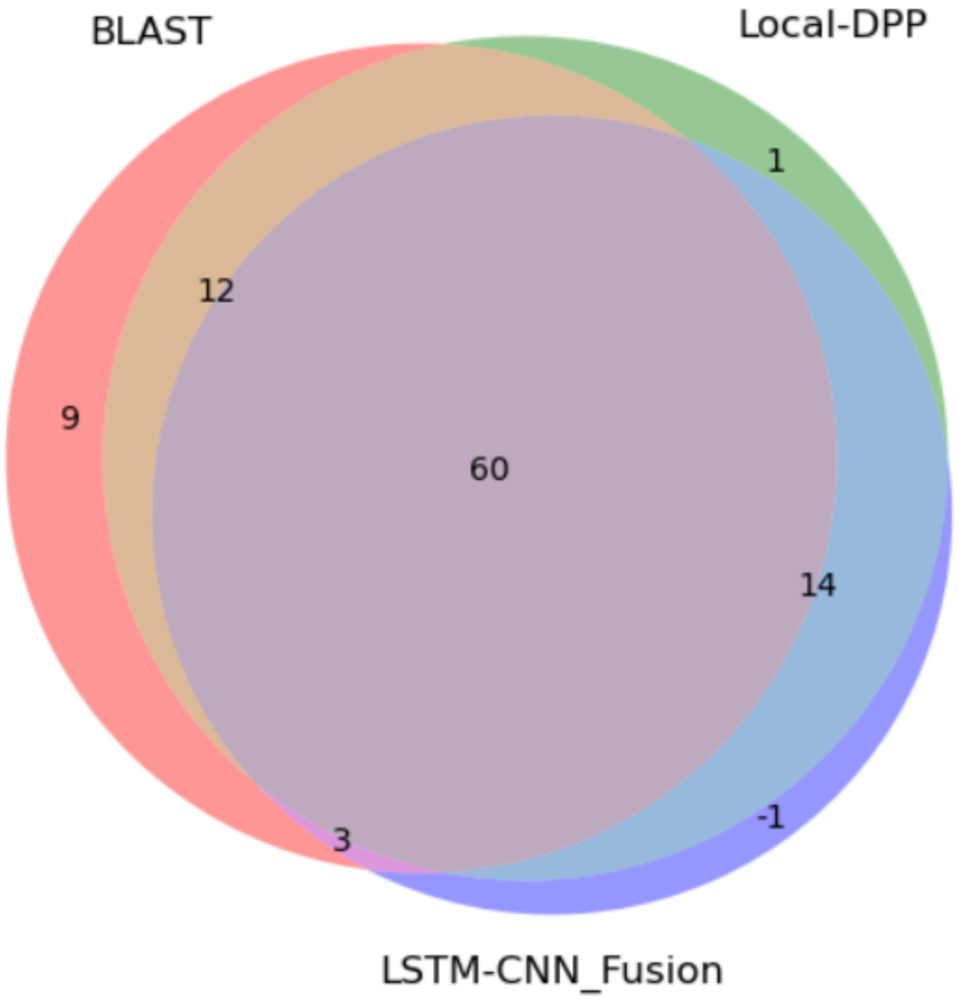


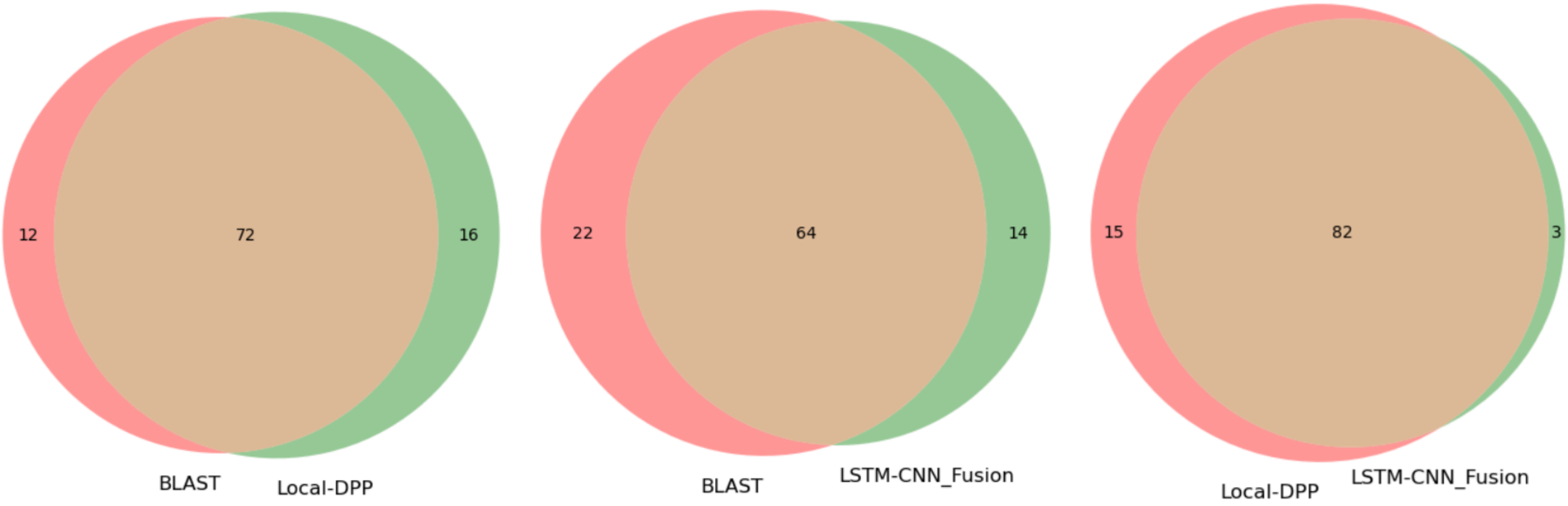

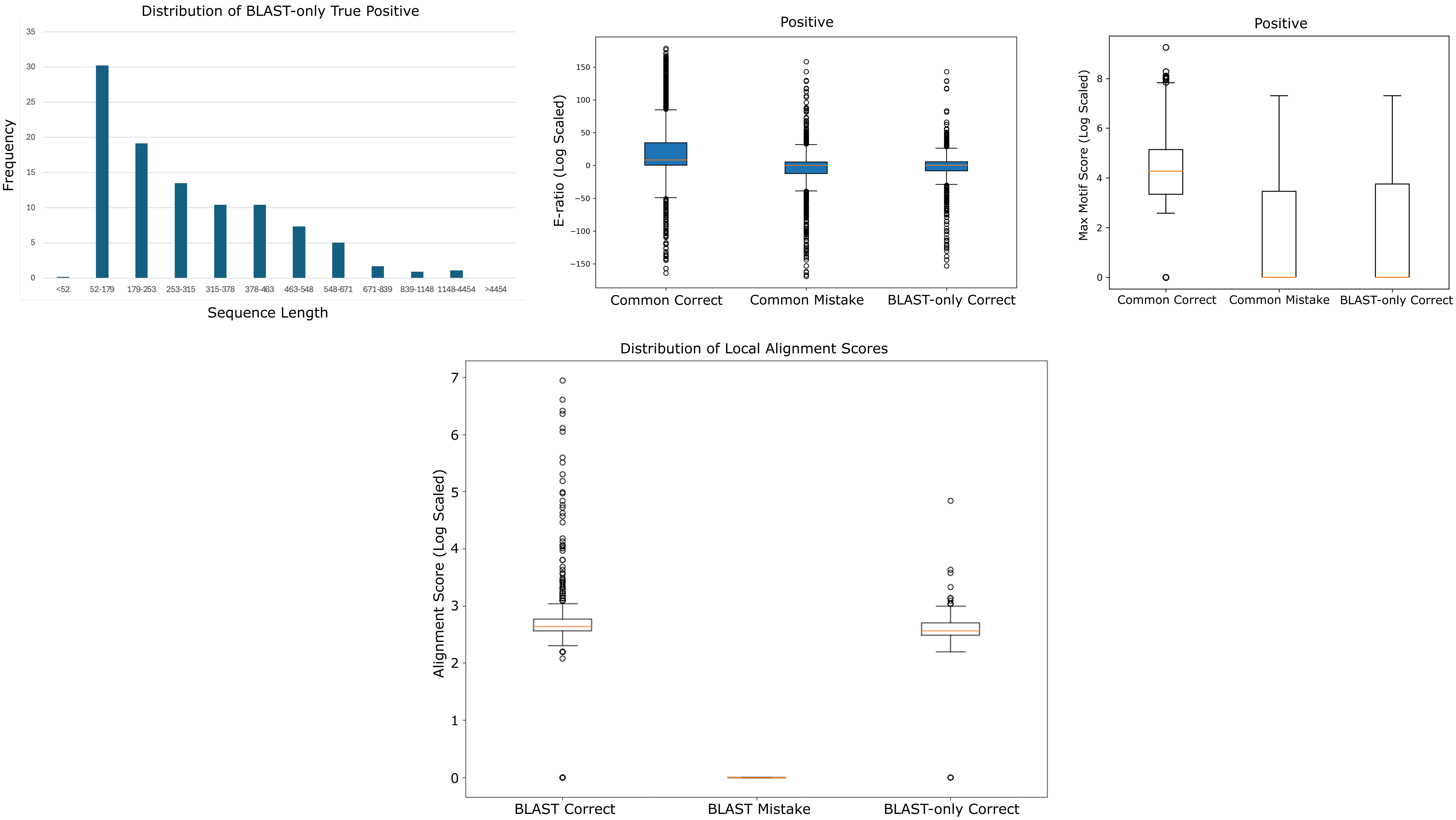


Fig S4: Analysis plots for DBP sequences correctly predicted by BLAST but mistaken by Local-DPP and LSTM-CNN_Fusion
