## Supplementary Data for "Benchmarking Recent Computational Tools for DNA-binding Protein Identification"

**Supplementary Data 1:**

Summary of replicability issues of the nine benchmarked tools

**Supplementary Data 2:**

Description of performance metrics used for evaluation

**Supplementary Data 3:**

ScanProsite performance on our proposed test set using different motif score thresholds

**Supplementary Data 4:**

Paper reported performance of the benchmarked nine tools

**Supplementary Data 5:**

Relative difference between paper reported sensitivity and EBTD & BTD sensitivity relative to reported sensitivity. We did not compare DNABP, because DNABP simply put away a small portion of its training set and used that as test set; we did not see data leakage problem in this test set.

**Supplementary Data 6:**

BLAST performance on proposed test set for different E-value thresholds

**Supplementary Data 7:**

Five-fold cross-validation performance of BLAST on BTD-Combo dataset

**Supplementary Data 8:**

Benchmarked tool summary with model and feature details

**Supplementary Data 9:**

HMMER performance on proposed test set for different E-value thresholds

**Supplementary Data 1:** Summary of replicability issues of the eleven benchmarked tools

| Tool | Issue |
|---|---|
| Local-DPP | None |
| DNABP | DNABR classifier software used for residue binding confidence score not currently available |
| iDNAProt-ES | feature index order not specified after feature selection |
| StackDPPred | Inconsistencies in feature extraction between code and paper |
| PseAAC | None |
| DeepDBP | Inconsistent feature count and model structure between code and paper |
| PDBP-Fusion | None |
| KK-DBP | feature index order not specified after feature selection |
| LSTM-CNN_Fusion | Some hyperparameters inconsistent between code and paper |
| PB_DBP | A lack of specified hyperparameters for the BiLSTM and final layers |
| PreDBP-PLMs | None |

**Supplementary Data 2:** Description of performance metrics used for evaluation

We used three performance metrics—sensitivity, specificity, and Matthews Correlation Coefficient (MCC)—to measure the tools' performance in each evaluation experiment. These metrics are defined as follows:

*Sensitivity* = $\frac{TP}{TP+FN}$

*Specificity* = $\frac{TN}{TN+FP}$

*MCC* = $\frac{(TP\times TN)-(FP\times FN)}{\sqrt{(TP+FP)\times(TP+FN)\times(TN+FP)\times(TN+FN)}}$

In these definitions, TP, FP, TN, and FN represent the number of true positives, false positives, true negatives and false negatives, respectively. Sensitivity and specificity range from 0 to 1, with 1 indicating a perfect score for positive and negative sample identification, respectively and 0 indicating the worst possible score. The MCC ranges from -1 to 1; where 1 denotes a perfect classifier, -1 denotes the worst classifier, and 0 indicates a random classifier. For MCC, the minority DNA-binding class (positive) samples have been labelled as positive in all cases.

One important point to note is that we did not include the conventional metric of accuracy. This decision was made because, in our proposed benchmarking dataset BTD, the negative class significantly outweighs the positive class. The dominance of one class would make accuracy a misleading metric for evaluating the effectiveness of the tools. Moreover, accuracy is a metric which depends on the ratio of positive and negative class sizes. Thus, the accuracy value reported on a test set can be very misleading if this ratio differs significantly from the ratio of real-world positive and negative population sizes.

**Supplementary Data 3:** ScanProsite performance on our proposed test set using different motif score thresholds

| SN | SP | MCC | Motif Score Threshold for Classification |
|---|---|---|---|
| 0.5806 | 0.7918 | 0.3723 | 0 |
| 0.4914 | 0.8468 | 0.3606 | 12.173 |
| 0.3739 | 0.8877 | 0.3085 | 18.1818 |
| 0.2521 | 0.9266 | 0.2479 | 26.9988 |
| 0.1267 | 0.9636 | 0.1709 | 41.1368 |

The four non-zero motif score thresholds are actually the 20th, 40th, 60th and 80th percentile of non-zero test set motif significance scores

**Supplementary Data 4:** Paper reported performance of the benchmarked eleven tools

| | | Paper Reported Performance | | |
|---|---|---|---|---|
| **Tool** | **Test Set** | **Sensitivity** | **Specificity** | **MCC** |
| Local-DPP | PDB186 | 0.925 | 65.6 | 0.625 |
| DNABP | PDB14K (406 seqs put away as test | 0.6847 | 0.7241 | 0.409 |
| iDNAProt-ES | PDB186 | 0.8131 | 0.8 | 0.613 |
| StackDPPred | PDB186 | 0.9247 | 0.8064 | 0.7363 |
| PseAAC | PDB186 | 0.95 | 0.688 | 0.666 |
| DeepDBP | PDB186 | 0.98 | 0.97 | 0.992 |
| PDBP-Fusion | PDB2272 | 0.7331 | 0.6685 | 0.5665 |
| KK-DBP | PDB186 | 0.978 | 0.645 | 0.661 |
| LSTM-CNN_Fusion | PDB2272 | 0.7623 | 0.9023 | 0.6463 |
| PB_DBP | Custom Dataset from Swiss-Prot | 0.975 | 0.945 | 0.92 |
| PreDBP-PLMs | PDB186 | 0.974 | 0.835 | 0.796 |
| | PDB2272 | 0.904 | 0.865 | 0.768 |

**Supplementary Data 5:** Relative difference between paper reported sensitivity and EBTD & BTD sensitivity relative to reported sensitivity.

| Tool | Sensitivity | | |
|---|---|---|---|
| | **Paper** | **EBTD** | **Deviation (%)** |
| Local-DPP | 0.925 | 0.97 | 4.6392 |
| StackDPPred | 0.9247 | 0.965 | 4.1762 |
| PDBP-Fusion | 0.7331 | 0.945 | 22.4233 |
| LSTM-CNN_Fusion | 0.7623 | 0.958 | 20.428 |

| | **Paper** | **BTD** | **Deviation (%)** |
|---|---|---|---|
| Local-DPP | 0.925 | 0.517 | -44.1081 |
| StackDPPred | 0.9247 | 0.427 | -53.8229 |
| PDBP-Fusion | 0.7331 | 0.499 | -31.9329 |
| LSTM-CNN_Fusion | 0.7623 | 0.502 | -34.1467 |

Note: We did not compare DNABP and PB_DBP. DNABP simply put away a small portion of its training set and used that as the test set. We did not see data leakage problem in this test set. The train and test set used by PB_DPB are not available and so, we do not know the exact limitations of their dataset.

**Supplementary Data 6:** BLAST performance on proposed test set for different E-value thresholds

| E-Value Threshold | SN | SP | MCC |
|---|---|---|---|
| 0.01 | 0.582346369 | 0.814653244 | 0.402012623 |
| 0.001 | 0.558882682 | 0.827181208 | 0.396611134 |
| 0.0001 | 0.545921788 | 0.836689038 | 0.397179968 |
| 0.00001 | 0.530502793 | 0.843736018 | 0.392515885 |
| 0.000001 | 0.517094972 | 0.85033557 | 0.389209565 |

**Supplementary Data 7:** Five-fold cross-validation performance of BLAST on BTD-Combo dataset

| | SN | SP | MCC |
|---|---|---|---|
| **Fold 1** | 0.576298 | 0.805612 | 0.385989 |
| **Fold 2** | 0.576298 | 0.805612 | 0.385989 |
| **Fold 3** | 0.58523 | 0.814794 | 0.40519 |
| **Fold 4** | 0.58324 | 0.809631 | 0.397124 |
| **Mean** | *0.5801* | *0.8097* | *0.3944* |
| **Std** | *0.004* | *0.0042* | *0.0083* |

**Supplementary Data 8:** Benchmarked tool summary with model and feature details

| Tool | Feature Type | Feature Detail | Computation Model | Model Detail |
|---|---|---|---|---|
| Local-DPP | Evolutionary | Local Pse-PSSM features | Classic | Random Forest |
| DNABP | Sequence + Evolutionary | PSSM with physicochemical properties (PSSM-PP),<br>Binding propensity measures (BP),<br>Non-binding propensity measures (NBP),<br>Physicochemical property feature (PHY) | Classic | Random Forest |
| iDNAProt-ES | Sequence + Evolutionary + Structure | Amino acid composition, Dubchak features,<br>PSSM Composition, PSSM Segmented Distribution<br>Secondary Structure Occurrence,<br>Secondary Structure Composition,<br>Accessible Surface Area Composition,<br>Torsional Angles Composition,<br>Structural Probabilities Composition,<br>Auto-Covariance (PSSM, Torsional Angles,<br>Structural Probabilities),<br>Bigram (PSSM, Torsional Angles, Structural Probablities) | Classic | SVM |
| StackDPPred | Evolutionary + Structure | PSSM-distance transformation (PSSM-DT) feature,<br>Residue probing transformation (RPT) feature,<br>Evolutionary distance transformation (EDT) feature,<br>Feature extracted from RCEM | Classic (stage) | SVM, Logistics Regression,<br>KNN, Random Forest |
| PseAAC | Sequence | Monogram, Bigram, Trigram, Gapped bigram,<br>Monogram percentile, Bigram percentile,<br>Nearest neighbor bigram | Classic | Extra Tree Classifier,<br>Random Forest |
| DeepDBP | Sequence | Same as features in PseAAC | Deep Learning | ANN, CNN |
| PDBP-Fusion | Sequence | One-hot encoding to DNA sequences | Deep Learning | CNN, LSTM |
| KK-DBP | Evolutionary | Reduced PSSM, PSSM-Composition, AADP-PSSM | Classic | Random Forest |
| LSTM-CNN_Fusion | Sequence + Evolutionary | One-hot encoding to DNA sequences, CNN to PSSM | Deep Learning | CNN, LSTM |
| PB_DBP | Sequence | ProtBert embedding of protein sequence | Deep Learning | PLM, BiLSTM |
| PreDBP-PLMs | Sequence + Evolutionary | ProtT5 embeddings<br>Pse-PSSM (Pseudo Position-Specific Score Matrix) | Deep Learning | PLM, CNN |

**Supplementary Data 9:** HMMER performance on proposed test set for different E-value thresholds

| E-value Threshold | SN | SP | MCC |
|---|---|---|---|
| 0.01 | 0.8536 | 0.571 | 0.4042 |
| 0.001 | 0.8346 | 0.6035 | 0.4144 |
| 0.0001 | 0.8346 | 0.6038 | 0.4146 |
| 0.00001 | 0.8344 | 0.6038 | 0.4144 |
| 0.000001 | 0.8344 | 0.6038 | 0.4144 |
